# PARP10 condensation inhibits viral infection via targeting NAD^+^ homeostasis

**DOI:** 10.64898/2025.12.22.696120

**Authors:** Xiaoxin Chen, Qi Xu, Yi Liu, Zhenshuo Zhu, Ziqiu Wang, Yun Zhang, Qi Chen, Le Tian, Zhiying Yao, Zhubing Shi, Peiguo Yang

## Abstract

ADP-ribosylation is a critical post-translational modification mediated by diphtheria toxin-like ADP-ribosyltransferases (ARTDs). The human ARTD family comprises 17 members that catalyze either poly- or mono-ADP-ribosylation. However, the functions and regulatory mechanisms of many ARTD proteins remain poorly understood. Here, we uncover an antiviral role for PARP10 through its ability to form biomolecular condensates. Viral infection triggers PARP10 condensation, a process driven by multivalent homotypic interactions among the structured domains. We show that mono-ADP-ribosylation of PARP10 suppresses its condensation, serving as a negative feedback mechanism that regulates condensate dynamics and protein stability. PARP10 condensates exhibit enhanced enzymatic activity, leading to decreased NAD^+^ levels and disruption of NAD^+^ homeostasis, ultimately inhibiting viral replication. Our findings establish PARP10 condensation as a novel mechanism for ARTD enzyme compartmentalization, with significant implications for innate immunity and host defense.

## Introduction

Cellular biochemistry is highly compartmentalized, with liquid-liquid phase separation (LLPS)-mediated biomolecular condensates serving as key regulatory hubs for diverse cellular processes^1,2^. Condensate formation modulates cellular functions through mechanisms such as spatial and temporal localization, enzymatic activation, buffering of signaling noise, and transcriptional regulation^1–3^. Numerous enzymes undergo phase separation to form condensates, where their enzymatic activity is often enhanced due to the increased local concentration^4,5^. However, enzymatic activity within condensates is also influenced by the local conformation and spatial arrangement of enzymes and substrates. Despite growing interest, the extent to which phase separation regulates enzymatic activity and the functional consequences of elevated activity within condensates remain poorly understood.

ADP-ribosylation is a post-translational modification involving the covalent attachment of ADP-ribose (ADPr) moieties to target proteins, using nicotinamide adenine dinucleotide (NAD^+^) as a cofactor^6–8^. This reaction is primarily catalyzed by the ADP-ribosyltransferases (ARTDs) or ‘ADPr writers’^7,9,10^. Following current nomenclature guidelines, we refer to these enzymes as ARTDs throughout this study^6^. Humans possess 17 ARTD family members, all of which contain a conserved ADPr transferase domain. ADP-ribosylation is reversible, with removal mediated by hydrolases, including macrodomain-containing hydrolases, poly(ADP-ribose) glycohydrolase (PARG), and ADP-ribosylhydrolase (ARH) enzymes^11,12^. PARP1 is best characterized for its role in DNA damage repair and maintenance of genome stability, and PARP1 inhibitors are clinically approved for cancer therapy^13–15^. Although all ARTD proteins are evolutionarily conserved in vertebrates and widely expressed across tissues, the biological functions of most ARTDs, particularly those that catalyze mono-ADP-ribosylation, remain largely uncharacterized^16^.

Recent studies on the evolutionary arms race between bacteria and bacteriophages have revealed NAD^+^ depletion as a widespread bacterial antiviral strategy. Bacteria encode various NAD^+^-hydrolyzing effector proteins that become activated upon phage infection, leading to rapid consumption of cellular NAD^+^. This metabolic collapse induces bacterial cell death and aborts phage propagation^17,18^. A similar mechanism has been identified in plant immunity, where NAD^+^ depletion or the generation of NAD^+^-derived secondary messengers activates downstream defense pathways^19,20^. In mammalian cells, the primary consumers of NAD^+^ include the ARTD and sirtuin (SIRT) families, SARM1, and extracellular NAD^+^ hydrolases such as CD38 and CD157^21,22^. SARM1, predominantly expressed in the nervous system, promotes axon degeneration upon activation^23,24^. Dysregulation of these NAD^+^-consuming enzymes and the consequent decline in NAD^+^ levels are also linked to aging and cellular senescence^21^. However, whether a coordinated, NAD^+^-centered antiviral mechanism operates in mammalian systems remains unclear.

In this study, we demonstrate that PARP10 undergoes phase separation to form biomolecular condensates during viral infection and plays a critical antiviral role in both human cell lines and mouse models. Using fluorescence microscopy, electron microscopy, and *in vitro* reconstitution assays, we establish that PARP10 condensation is driven by multivalent homotypic interactions involving structured domains, particularly the N-terminal RNA recognition motifs (RRMs). We further propose a negative feedback loop in which auto-ADP-ribosylation regulates the dynamics and stability of PARP10 condensates. Both the ADP-ribosyltransferase activity and the capacity for phase separation are essential for antiviral function of PARP10. Mechanistically, PARP10 condensation enhances its enzymatic activity, leading to substantial depletion of cellular NAD^+^ and disruption of NAD^+^ homeostasis, thereby inhibiting viral replication. Together, our findings uncover a novel role for PARP10-driven condensates in antiviral immunity through targeted modulation of NAD^+^ metabolism.

## Results

### PARP10 is an antiviral factor

We performed RNA-seq analysis on vesicular stomatitis virus (VSV)-infected bone marrow-derived macrophages (BMDMs). The upregulated genes induced by VSV largely overlapped with previously identified interferon-stimulated genes (ISGs)^25^ and were enriched in immune-related pathways, confirming the validity of our cellular model (Extended Data Fig. 1a-c). Among the differentially expressed genes, eight *ARTD* genes were significantly upregulated (*Parp11/10/12/3/9/14/8/7*), and two were significantly downregulated (*Parp1/16*) (Fig. 1a and Supplementary Table 3). To assess the impact of ARTD proteins on VSV replication, we generated expression constructs for 15 human *ARTD* genes, excluding PARP2 and PARP4 due to their large coding sequences. Ectopic expression of individual ARTDs in HEK293T cells revealed that PARP10, PARP12, and PARP9 strongly inhibited VSV replication, with PARP10 exhibiting the most potent antiviral effect (Fig. 1b). The remaining ARTDs showed no significant effect, with PARP1 and PARP6 modestly enhancing viral replication (Fig. 1b). Given its robust activity, we focused subsequent analyses on PARP10.

**Fig. 1.**
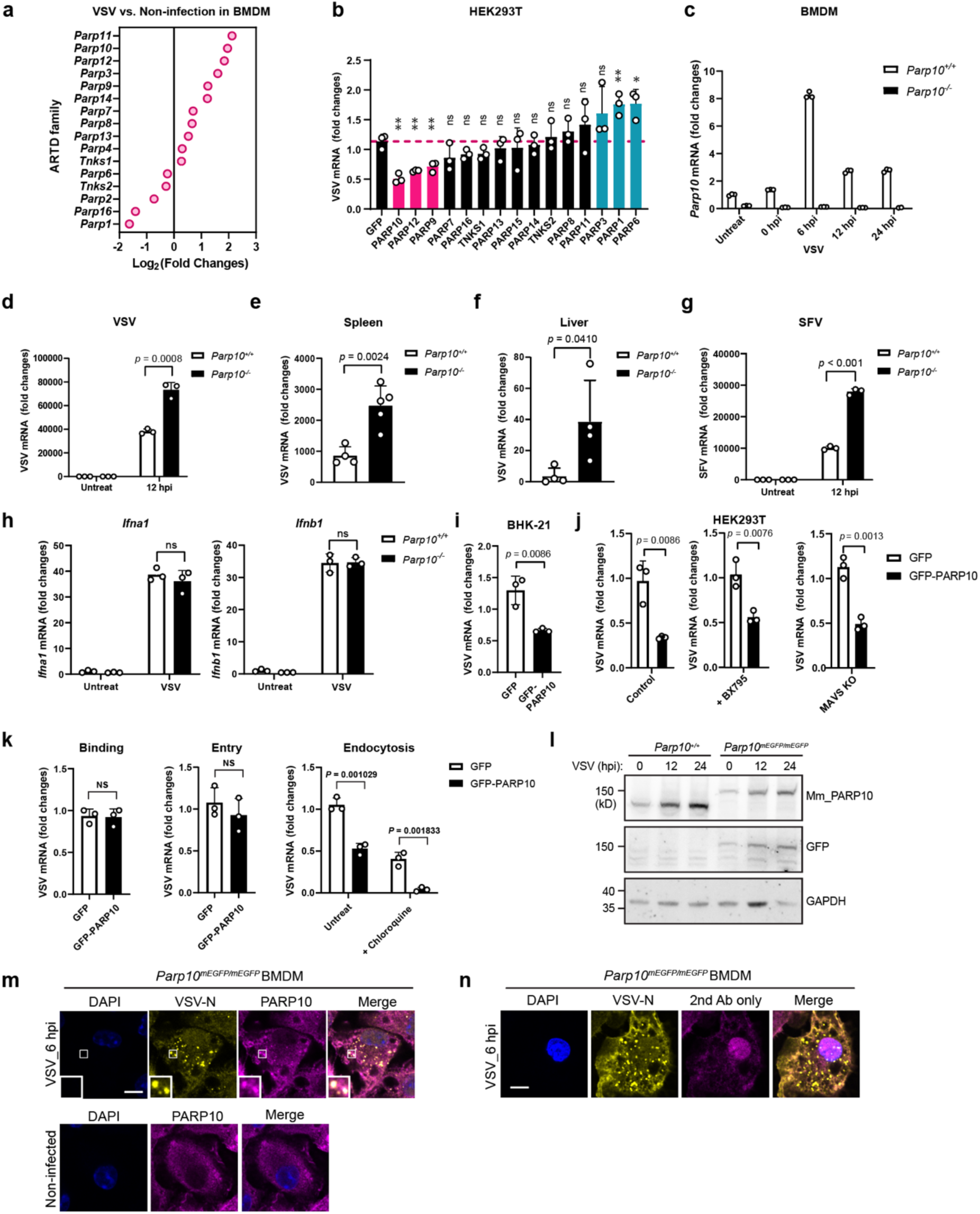
PARP10 is an antiviral factor. **a**, Fold changes of differentially expressed *ARTD* genes in BMDM with VSV (12 hpi, MOI = 1.5) infection compared to non-infected control. *n* = 3. **b**, Normalized VSV mRNA levels in HEK293T cells transfected with different ARTD genes and infected with VSV (8 hpi, MOI = 1.5). *n* = 3. **c**, Normalized PARP10 mRNA levels in *Parp10^+/+^* and *Parp10^-/-^*BMDM at different time points post-VSV (MOI = 1.5) infection. *n* = 3. **d**, Normalized VSV mRNA levels in *Parp10^+/+^* and *Parp10^-/-^* BMDM infected with VSV (12 hpi, MOI = 1.5). *n* = 3. **e**,**f**, Normalized VSV mRNA levels in spleen (**e**) and liver (**f**) samples isolated from VSV (1x10^8^ PFU/mouse, 24 hpi) infected *Parp10^+/+^* (*n* = 4) and *Parp10^-/-^* (*n* = 5) mice. **g**, Normalized SFV mRNA levels in *Parp10^+/+^* and *Parp10^-/-^* BMDM infected with SFV (12 hpi, MOI = 0.5). *n* = 3. **h**, Normalized *Ifna1* and *Ifnb1* mRNA levels in *Parp10^+/+^* and *Parp10^-/-^* BMDM infected with VSV (12 hpi, MOI = 1.5). *n* = 3. **i**, Normalized VSV mRNA levels in BHK-21 cells transfected with GFP or GFP-PARP10 and infected with VSV (8 hpi, MOI = 1.5). *n* = 3. **j**, Normalized VSV mRNA levels in WT/BX795-treated/MAVS KO HEK293T cells transfected with GFP or GFP-PARP10, followed by VSV infection (8 hpi, MOI = 1.5). *n* = 3. **k**, Normalized VSV mRNA levels in HEK293T cells transfected with GFP or GFP-PARP10 during viral binding, entry, and endocytosis steps. The endocytosis steps were blocked with 10 μM Chloroquine. **l**, Western blot showing the expression of mEGFP-PARP10 in *Parp10^+/+^* and *Parp10^mEGFP/mEGFP^* BMDM with or without VSV infection (MOI = 1.5) for the indicated times. **m**, Representative images of colocalization between endogenous PARP10 and VSV inclusion labeled by N protein in *Parp10^mEGFP/mEGFP^* BMDM infected with VSV (6 hpi, MOI = 1.5) or in non-infection control. **n**, Control IF experiment as (**m**) except without primary antibody staining for PARP10, thus excluding the leaky signal from VSV-N channel. Data are shown as mean ± SD. in (**b**)-(**k**). Scale bar: 10 μm for (**m**) and (**n**).

Although PARP10 has been implicated in tumorigenesis^26–28^, its role in antiviral defense remains poorly understood. The amino acid sequence alignment revealed over 60% conservation between human and mouse PARP10 (Extended Data Fig. 1d). To investigate its antiviral function, we generated a *Parp10* knockout (KO) mouse model using the CRISPR-Cas system. Double gRNAs mediated complete deletion of *Parp10* coding region (Extended Data Fig. 1e). Western blot confirmed loss of PARP10 protein expression in tissues from *Parp10* KO mice (Extended Data Fig. 1f). PARP10 exhibited broad tissue expression, with higher levels in immune-related organs such as bone marrow, lymph nodes, spleen, and lung (Extended Data Fig. 1f,g). Single-cell RNA-seq data from Human Protein Atlas indicate that human PARP10 is ubiquitously expressed across tissues, with prominent expression in immune cells, including dendritic cells, natural killer (NK) cells, and T-cells (Extended Data Fig. 1h). RT-qPCR analysis showed that *Parp10* expression was induced upon VSV infection and was undetectable in BMDMs from *Parp10* KO mice (Fig. 1c).

To assess the role of PARP10 in VSV replication, we isolated BMDMs from wild-type (WT) and *Parp10* KO mice and infected the cells with VSV. RT-qPCR analysis showed significantly enhanced VSV replication in *Parp10* KO BMDMs compared to WT controls (Fig. 1d). Similarly, VSV replication was markedly increased in spleen and liver tissues from *Parp10* KO mice relative to WT mice following infection (Fig. 1e,f). Furthermore, we demonstrated that PARP10 restricts not only the negative-stranded RNA virus VSV but also the positive-stranded RNA virus Semliki Forest virus (SFV) (Fig. 1g). Collectively, these findings establish PARP10 as an antiviral factor, functioning both *in vitro* and *in vivo*.

### The antiviral activity of PARP10 is independent of the innate immune signaling pathway

To evaluate a potential role of PARP10 in the innate immune response, particularly in the type I interferon (IFN) signaling pathway, we assessed the expression of type I interferon genes. Upon VSV infection, induction of *Ifna1* and *Ifnb1* was comparable between *Parp10* KO and WT BMDMs (Fig. 1h). Furthermore, PARP10 retained strong antiviral activity against VSV in BHK-21 cells (Fig. 1i), which are defective in interferon signaling, suggesting an interferon-independent mechanism. To further investigate this, we pharmacologically inhibited TANK-binding kinase 1 (TBK1), a central mediator of type I IFN signaling, using BX795. PARP10-mediated suppression of viral replication remained largely unaffected by BX795 treatment (Fig. 1j). This finding was confirmed in MAVS-knockout 293T cells, where PARP10 still effectively inhibited VSV replication despite the absence of MAVS (Fig. 1j and Extended Data Fig. 1i). The efficacy of BX795 treatment and MAVS KO was validated by impaired expression of downstream interferon-stimulated genes (Extended Data Fig. 1j). Additionally, we examined the phosphorylation of interferon regulatory factor 3 (IRF3), a hallmark of type I IFN pathway activation, and found that VSV-induced IRF3 phosphorylation was unaltered in *Parp10* KO BMDMs (Extended Data Fig. 1k). These results collectively indicate that PARP10 does not modulate the canonical MAVS-IRF3-dependent IFN signaling pathway and that its anti-VSV effect is not mediated through enhanced innate immune activation. Time-course analysis of viral replication revealed no differences in VSV binding, entry, or endocytosis in PARP10-expressing cells (Fig. 1k), further supporting a direct or non-immune-mediated antiviral mechanism.

### PARP10 forms subcellular foci during viral infection

To examine the subcellular localization of PARP10 during VSV infection, we generated mEGFP knock-in (KI) mice by inserting the mEGFP coding sequence in-frame at the N-terminus of the mouse *Parp10* gene (Extended Data Fig. 1l). Western blot analysis of BMDMs revealed comparable PARP10 protein expression levels in WT and *mEGFP-Parp10* KI BMDMs (Fig. 1l). VSV infection induced PARP10 expression, indicating that the KI allele is functionally responsive (Fig. 1l). In uninfected BMDMs, mEGFP-PARP10 exhibited a diffuse cytoplasmic distribution, whereas upon VSV infection, it relocalized to distinct foci (Fig. 1m). These PARP10 foci colocalized with VSV inclusions labeled by the viral nucleocapsid (N) protein (Fig. 1m). Due to the relatively weak native GFP signal, we enhanced detection using anti-GFP immunostaining. Control experiments omitting the GFP antibody showed no detectable foci, ruling out bleed-through from the VSV-N channel and confirming the specificity of the observed PARP10 foci (Fig. 1n). To further characterize PARP10 foci in human cells, we employed the human osteosarcoma cell line U2OS, which possesses a large cytoplasmic volume conducive to high-resolution subcellular imaging. In infected U2OS cells, PARP10 foci were observed within VSV inclusions (Fig. 2a), indicating that PARP10 foci formation is a conserved response to viral infection across mouse and human systems.

**Fig. 2.**
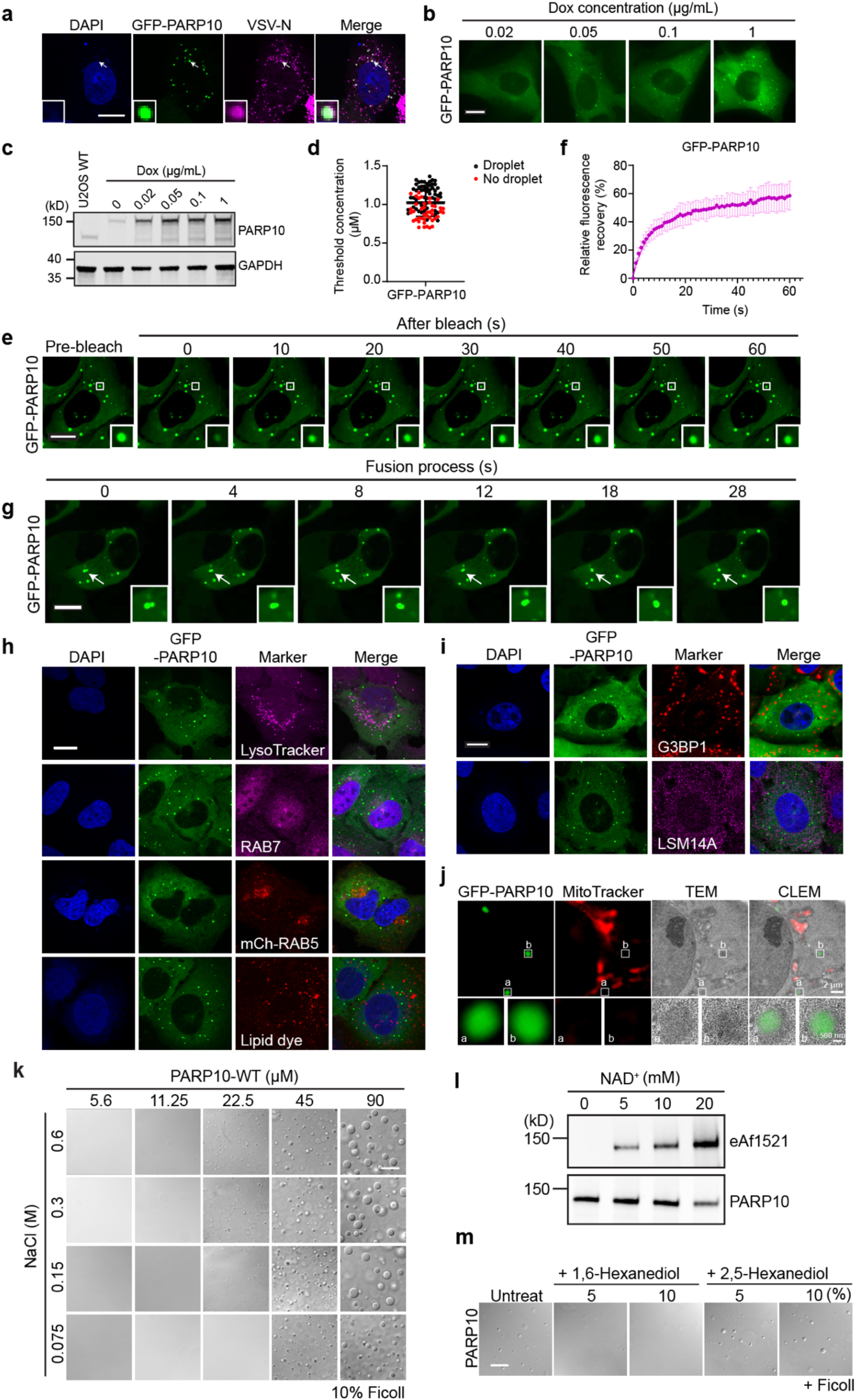
PARP10 forms condensates in cells and *in vitro*. **a**, Representative images to show the colocalization between GFP-PARP10 foci and VSV inclusions. **b**, Representative images of GFP-PARP10 inducible U2OS cells stimulated with different doxycycline doses. Higher expression induced more foci formation. **c**, Western blot showing the protein expression of GFP-PARP10 with different doxycycline doses. **d**, Calculation of the cellular threshold concentration of GFP-PARP10 to form cytoplasmic foci. The absolute concentration in cells is converted by a standard curve created by measuring the intensity of recombinant PARP10 protein. Each black dot indicates one cell with condensates, the red dot indicates the cell without condensates, and the line marks the threshold concentration for condensate formation. At least 100 cells were measured and analyzed. **e**,**f**, FRAP experiments for GFP-PARP10 foci in cells. The fluorescent intensity in the region of interest (ROI) in (**e**) is quantified in (**f**). **g**, Representative images for the GFP-PARP10 droplets fusion process. **h**, Representative images of GFP-PARP10 foci with LysoTracker, lipid dye, RAB7 staining, and mCherry-RAB5 co-expression. **i**, Representative images for GFP-PARP10 foci in cells stained with G3BP1 and LSM14A. **j**, Correlative light electron microscopy (CLEM) images of GFP-PARP10 foci. GFP-PARP10 foci were identified by the high electron density and correlation with GFP fluorescence. MitoTracker in red indicates the location of mitochondria. **k**, Representative images of recombinant PARP10 protein condensation induced by 10% Ficoll under the indicated salt and protein concentrations. **l**, Western blot showing the ADP-ribosylation of recombinant PARP10 with the indicated concentrations of NAD^+^ *in vitro*. The ADPr-modified PARP10 was blotted with eAf1521. **m**, Representative images of Ficoll-induced PARP10 condensation *in vitro* with the treatment of 1,6-Hexanediol or 2,5-Hexanediol. Data are shown as mean ± SD. in (**d**) and (**f**). Scale bar: 10 μm for (**a**), (**b**), (**e**), (**g**)-(**i**). 2 μm for (**j**) and 500 nm for the inset. 20 μm for (**k**) and (**m**).

### PARP10 foci are membrane-less biomolecular condensates

To assess the capacity of PARP10 to form subcellular foci, we established a doxycycline-inducible GFP-PARP10 expression system in U2OS cells on a PARP10 KO background. PARP10 KO was confirmed by western blotting and immunofluorescence staining using both in-house-generated and commercially available PARP10-specific antibodies (Extended Data Fig. 2a-c). We observed a doxycycline dose-dependent formation of cytoplasmic PARP10 foci (Fig. 2b,c), indicating that PARP10 foci formation is an autonomous process rather than recruitment into pre-existing cellular structures. Using recombinant protein to generate a standard curve, we estimated the threshold concentration for GFP-PARP10 foci formation to be appropriately 1 μM (Fig. 2d). PARP10 expression was induced by IFNβ and IFNγ, but not by TNFα (Extended Data Fig. 2d), supporting its classification as a bona fide ISG. Fluorescence recovery after photobleaching (FRAP) assay indicated that PARP10 compartments were dynamic and exhibited liquid-like features (Fig. 2e,f). Time-lapse live-cell imaging captured fusion events between PARP10 foci (Fig. 2g). These foci did not co-localize with membrane-associated markers, including RAB5, RAB7, lysotracker, and lipid dye-stained vesicles (Fig. 2h), nor with markers of stress granules (G3BP1) or P-bodies (LSM14A) (Fig. 2i). Correlative light and electron microscope (CLEM) further confirmed that PARP10 foci are distinct, membrane-free structures (Fig. 2j). Collectively, these findings indicate that PARP10 forms membraneless organelles that fulfill the criteria for biomolecular condensates.

We also reconstituted PARP10 phase separation *in vitro* using recombinant protein purified from a bacterial expression system. PARP10 underwent phase separation in the presence of 10% Ficoll as a molecular crowding agent (Fig. 2k). This phase separation was insensitive to NaCl concentrations ranging from 0.075 M to 0.6 M, indicating robustness across physiological salt conditions (Fig. 2k). In the presence of NAD^+^, recombinant PARP10 underwent auto-ADP-ribosylation (Fig. 2l). Treatment with 1,6-hexanediol, but not its structural isomer 2,5-hexanediol, disrupted PARP10 condensates, suggesting that hydrophobic interactions play a role in its condensation (Fig. 2m). The *in vitro* reconstitution assay demonstrates the intrinsic ability of PARP10 to undergo phase separation.

### Multivalent interactions mediate PARP10 condensation

PARP10 contains three RNA recognition motifs (RRMs, RRM1/2/3) at its N-terminus, two ubiquitin-interacting motifs (UIMs) adjacent to the C-terminal catalytic domain, and two centrally located K Homology (KH) domains, KH1 and KH2 (Fig. 3a)^29^. To dissect the molecular basis of PARP10 condensation, we generated a series of deletion mutants targeting predicted structured regions (Fig. 3b). To assess ADP-ribosylation activity, we employed the well-characterized eAf1521-rFC reagent from *Archaeoglobus fulgidus,* which recognizes both mono-ADPr and various poly-ADPr chains^30,31^. Basal cytoplasmic ADPr signal was primarily mitochondrial in origin (Extended Data Fig. 3a), consistent with prior reports^32^. PARP10 expression enhanced the cellular ADPr signal, whereas catalytic deletion (ΔCD) abolished this signal and revealed the underlying basal mitochondrial signal (Fig. 3c), confirming proper expression and functionality of the constructs in cells. Among all the constructs tested, deletion of either the RRM or KH domains nearly abolished condensate formation (Fig. 3c,d). Similarly, removal of the UIMs significantly impaired condensation, despite comparable protein expression levels (Fig. 3c-e).

**Fig. 3.**
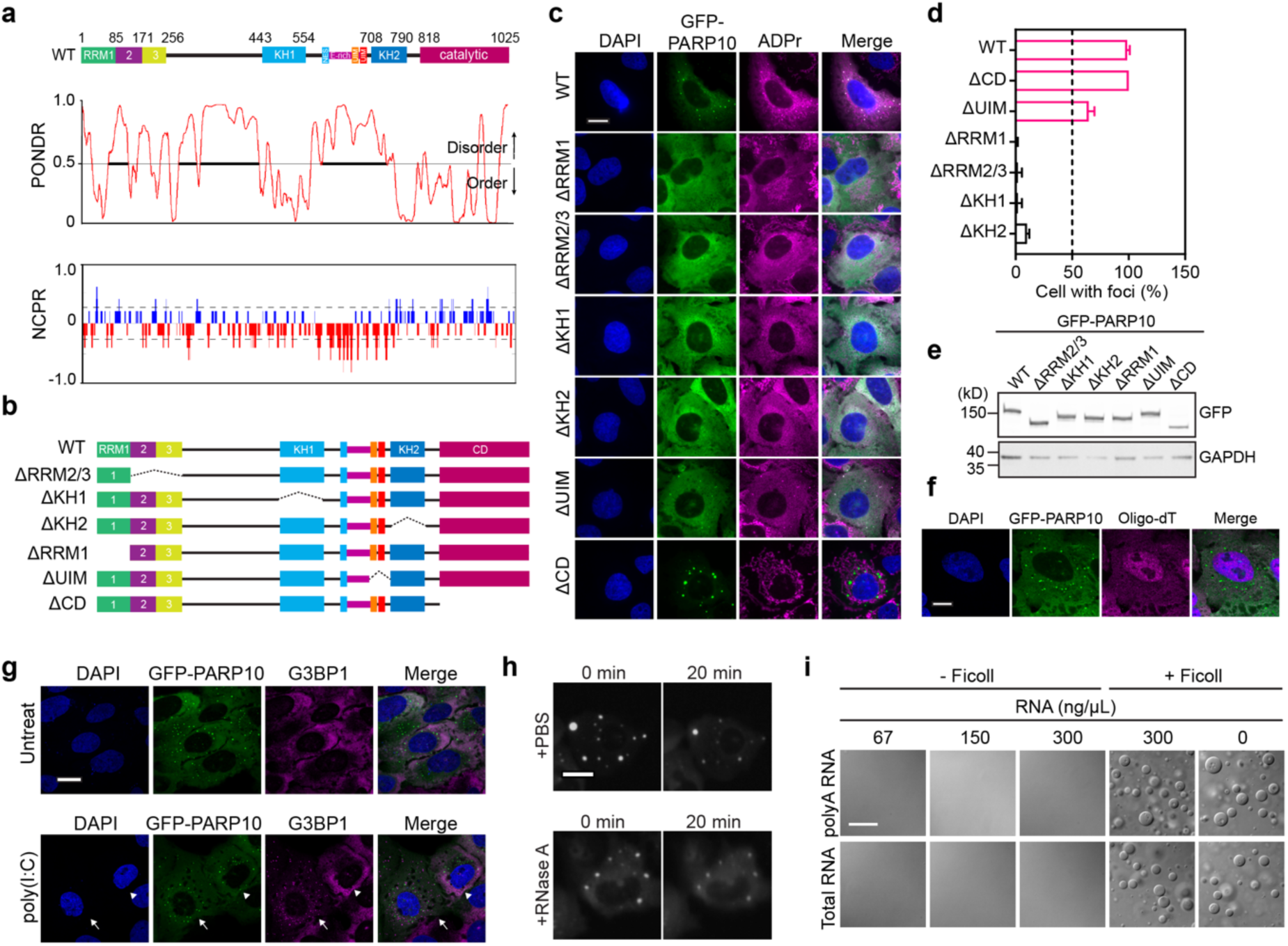
Multivalent interactions across PARP10 contribute to condensation. **a**, The schematic diagram for PARP10 protein and predicted intrinsically disordered region (IDR) and net charge per residue (NCPR) pattern are shown below. **b**, The schematic diagram for PARP10 mutants used. **c**, Representative images of cells stably expressing PARP10 mutants in (**b**). **d**, Quantification of the percentage of cells with foci in (**c**). **e**, Western blot showing the relative protein expression of each construct used in (**c**). **f**, Representative images of oligo-dT FISH staining in GFP-PARP10 condensate. **g**, Representative images of GFP-PARP10 condensates with poly(I:C) treatment. Cells with G3BP1 foci were indicative of successful poly(I:C) transfection. Arrow indicates cells transfected with poly(I:C) and arrowhead indicates cells absent of poly(I:C). **h**, Representative images of GFP-PARP10 condensates with RNase A (100 μg/mL) treatment for 20 min. PBS was used as control buffer. **i**, Representative images of PARP10 WT condensation in the presence of indicated concentrations of polyA RNA or total RNA with or without 10% Ficoll. Data are shown as mean ± SD. in (**d**). Scale bar: 10 μm for (**c**), (**f**)-(**h**). 20 μm for (**i**).

The RRM domains were initially identified based on sequence homology with canonical RRMs. Sequence alignment with RRMs from hnRNPA1 and TDP43 revealed overall similarity (Extended Data Fig. 3b). However, PARP10 RRM1 lacks the conserved phenylalanine residue critical for RNA binding in these proteins (Extended Data Fig. 3b). Moreover, there is currently no experimental evidence supporting direct RNA binding by PARP10 RRMs. Although PARP10 has been reported to ADP-ribosylate phosphorylated RNA termini^33^, it remains unclear whether the RRMs facilitate RNA substrate recognition. Given the critical role of RRMs in condensation, we next examined whether RNA is a component of PARP10 condensates. Oligo-dT fluorescence in situ hybridization (FISH) revealed no detectable poly(A) RNA within PARP10 foci (Fig. 3f). To further assess RNA dependence, we utilized poly(I:C), a synthetic double-stranded RNA analog that activates RNase L and induces global degradation of cytoplasmic RNA^34^. In poly(I:C)-transfected cells exhibiting G3BP1-positive stress granules, indicative of a responsive cellular state, PARP10 condensates remained unchanged (Fig. 3g). Additionally, treatment with RNase A in permeabilized cells did not dissolve PARP10 droplets (Fig. 3h). *In vitro*, neither the addition of total RNA nor poly(A) RNA induced PARP10 condensation, nor did they alter Ficoll-induced phase separation (Fig. 3i). These results collectively indicate that RNA is neither a structural scaffold nor a required modulator of PARP10 condensate formation under the conditions tested.

Expression of individual PARP10 fragments in cells revealed that constructs containing both RRM and KH domains formed cytoplasmic foci, whereas other fragments exhibited diffuse localization (Extended Data Fig. 3c,d), suggesting that RRM and KH folded domains facilitate self-association. Consistent with this, co-immunoprecipitation assays demonstrated that the RRM and KH domains engage in both homotypic and heterotypic interactions (Extended Data Fig. 3e). These findings indicate that multivalent interactions mediated by structured domains are essential for efficient PARP10 condensation.

### N-terminal RRM domains mediate PARP10 oligomerization and phase separation

We further examined the self-association behavior of recombinant PARP10 using single-particle cryogenic electron microscopy (cryo-EM). 2D classification revealed both open and closed ring-like structures of variable sizes (Fig. 4a and Extended Data Fig. 4a,b). Open rings typically consist of more than five triangular protomers with diameters ranging from ∼155 Å to 200 Å and inter-protomer angles of appropriately 120-155°. A closed ring composed of nine protomers was observed, exhibiting an internal angle of ∼140°. Dimers of triangular protomers were also detected and represent the minimal assembly unit of the ring structure. 3D reconstruction and refinement yielded three medium-resolution cryo-EM maps, each containing four to six discernible protomers (Fig. 4b and Extended Data Fig. 4c). The protomers within a ring are not coplanar; instead, they twist relative to each other, forming a helical arrangement (Extended Data Fig. 4d), a configuration that may promote the formation of extended interaction networks.

**Fig. 4.**
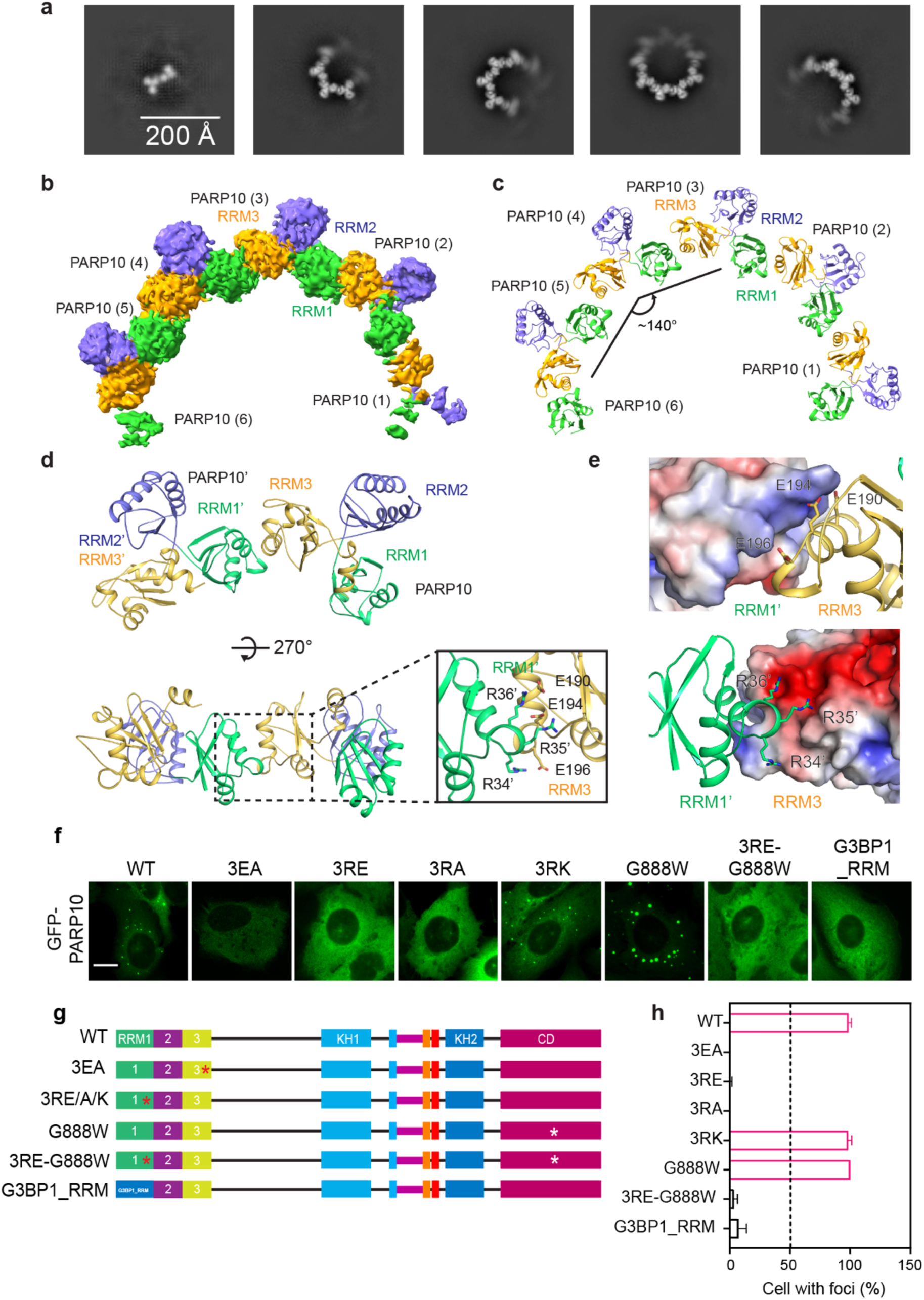
N-terminal RRM domains mediate the oligomerization of PARP10. **a**, Representative 2D classes of the recombinant, full-length PARP10 protein under cryo-EM. Scale bar: 200 Å. **b**, Cryo-EM map of PARP10 RRM1-3 pentamer obtained from 3D reconstruction and refinement. **c**, Structural model of PARP10 RRM1-3 pentamer, which is generated by docking AlphaFold-predicted models of PARP10 RRM1-3 domains into the cryo-EM map shown in (**b**). The PARP10 RRM1-3 oligomer adopts an open ring conformation with an angle of 140 degrees. **d**, Detailed illustration of the RRM1-3 dimer interface at two orientations. One dimeric interface of RRM1-3 is zoomed in, and residues in the interface between two RRM1-3 protomers are shown in sticks. **e**, Two protomers of PARP10 RRM1-3 contact with each other by electrostatic interactions. One protomer is represented with electrostatic potential surface, and the other is shown in cartoon. **f,g**, Representative images of cells (**f**) stably expressing PARP10 mutants depicted in (**g**). Scale bar: 10 μm. **h**, Quantification of the percentage of cells with foci in (**f**).

Further 3D classification, focusing on particles with the highest number of triangular units, enabled the generation of a refined cryo-EM map for structural analysis (Extended Data Fig. 4e-h). PARP10 contains three N-terminal domains structurally homologous to RRMs^29^. Guided by AlphaFold3-predicted models, we fitted the N-terminal RRM1-3 domains into the cryo-EM density, building a hexameric model of the RRM1-3 region. Clear density was observed for four of the six subunits, while the remaining two were partially resolved (Fig. 4b,c). Other regions of PARP10, including the catalytic domain, were not visible, likely due to intrinsic flexibility and the small size of the intervening intrinsically disordered region (IDR) and C-terminal segment.

In the structural model, the RRM1-3 domains adopt a triangular conformation. Ring assembly occurs through a head-to-tail interaction between RRM1 of one protomer and RRM3 of the adjacent protomer. At this interface, residues R34, R35, and R36 of RRM1 are positioned to interact with a negatively charged patch on RRM3, primarily through electrostatic interactions with acidic residues E190, E194, and E196 (Fig. 4d,e). To assess the functional importance of these residues at the RRM1/3 interaction interface, we substituted three key glutamate residues with alanine (3EA) in RRM3 and three arginine residues with alanine (A) or glutamate (E) to neutralize or reverse their charge in RRM1. All mutations abolished condensate formation, phenocopying RRM deletion (Fig. 4f-h). In contrast, a 3RK mutant, which preserves the positive charge, retained the ability to form condensates, underscoring the importance of electrostatic interactions in this process. The 3R motif was also essential for condensation of the G888W mutant, which lacks catalytic activity (Fig. 4f-h). Furthermore, replacing PARP10 RRM1 with the canonical RRM of G3BP1 failed to restore condensation, indicating that the specific structural or biophysical properties of PARP10 RRM1 are critical (Fig. 4f-h).

Notably, rare higher-order architectures were observed in 2D class averages (Extended Data Fig. 4i,j). One class displayed an S-shaped configuration, in which the direction of the ring path is altered to reverse direction in the middle (Extended Data Fig. 4i). In another class, two rings are interconnected by four copies of the third domain (RRM3) that are not involved in forming each ring structure, resulting in a rhomboid shape at the contact region and forming a linked ring structure (Extended Data Fig. 4j). These higher-order structures, together with the spiral arrangement of the ring, likely expand the diversity of PARP10 self-association, thereby facilitating PARP10 oligomerization and phase separation.

### PARP10 condensates exhibit catalytic activity

To determine whether PARP10 condensates are catalytically active, we performed immunofluorescence staining using ADP-ribose (ADPr)-specific antibodies. The commercial 10H antibody detects ADPr chains containing more than ten units, whereas the E6F6A antibody recognizes both mono- and poly-ADP-ribosylation. Immunofluorescence analysis revealed that PARP10 condensates were enriched with ADPr signals as detected by E6F6A but showed minimal 10H staining, indicating the absence of long poly-ADPr chains within these condensates (Fig. 5a). The presence of mono-ADPr was further confirmed using the Macro2-3 domain of human PARP14 (Fig. 5b), a well-established reader domain specific for mono-ADP-ribosylation^35^. The G888W mutation in the catalytic triad abolishes its enzymatic activity^36^, and no ADPr signal was observed in condensates formed by this mutant (Fig. 5b). Notably, PARP10 G888W still formed condensates, demonstrating that the catalytic activity is dispensable for condensate formation.

**Fig. 5.**
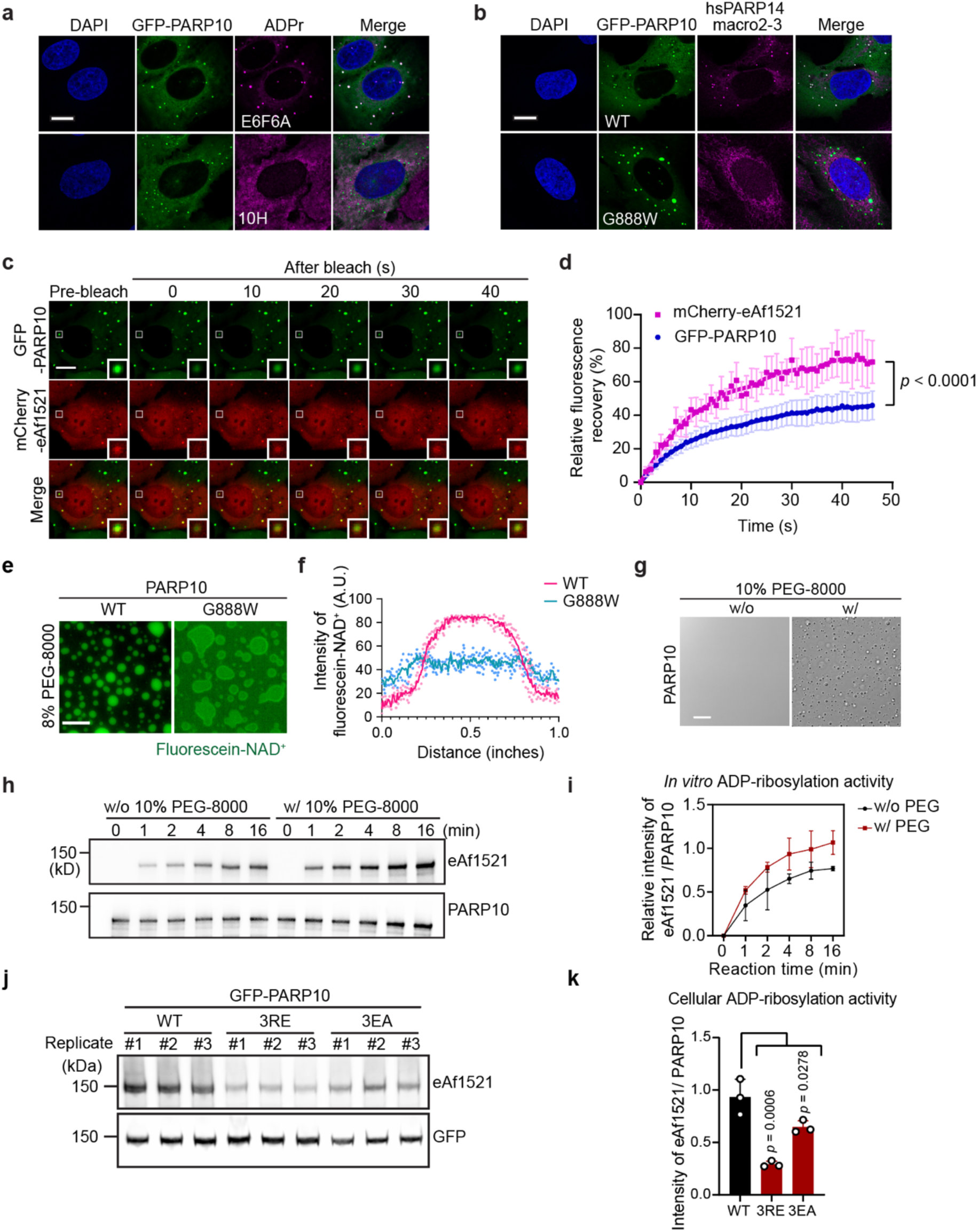
PARP10 condensation promotes ADP-ribosylation activity. **a**,**b**, Representative images for GFP-PARP10 condensate in cells stained with ADPr recognition antibodies (E6F6A and 10H) (**a**) and reagent (hsPARP14 macro2-3-rFC) (**b**). The catalytic deficient mutant GFP-PARP10 G888W served as a negative control. **c**,**d**, FRAP experiments for GFP-PARP10 and mCherry-eAf1521 co-expression cells. The fluorescent intensity in the region of interest (ROI) in (**c**) is quantified in (**d**). **e**,**f**, Representative images (**e**) and line quantification (**f**) of fluorescein-NAD^+^ enrichment in PARP10-WT but not G888W condensates induced by PEG-8000 *in vitro*. **g-i**, Representative images of the reactions to show the PEG-8000 induced PARP10 condensation (**g**). Western blot showing ADP-ribosylation of 10 μM PARP10 with or without 10% PEG-8000 addition (**h**). The quantification of the ADP-ribosylation activity was shown in (**i**). *n* = 2. **j**,**k**, Western blot showing the ADP-ribosylation activity of PARP10 WT and condensation-deficient mutants 3RE and 3EA in cells (**j**). ADP-ribosylation levels were blotted with eAf1521. The quantification of the ADP-ribosylation activity was shown in (**k**). *n* = 3. Data are shown as mean ± SD. in (**d**), (**i**), (**k**). Scale bar: 10 μm for (**a**)-(**c**); 20 μm for (**e**) and (**g**).

To further assess ADPr dynamics, we employed mCherry-eAf1521, a cellular ADPr biosensor. Under normal growth conditions, this reporter exhibited diffuse localization in U2OS cells. Upon co-expression with WT PARP10, mCherry-eAf1521 accumulated within PARP10 condensates, confirming the presence of ADPr in these structures (Fig. 5c). Fluorescence recovery after photobleaching (FRAP) analysis of ADPr signals within PARP10 condensates revealed rapid exchange between ADPr-modified substrates and the surrounding cytoplasm, indicating dynamic substrate shuttling in and out of the condensates (Fig. 5c,d). The mobility of ADPr signals exceeded that of PARP10 itself, suggesting that the condensates function as catalytic hubs facilitating the modification of diverse substrates.

In the *in vitro* reconstituted PARP10 condensation assay, droplets formed by WT PARP10, but not the catalytically inactive G888W mutant, exhibited strong enrichment of the cofactor fluorescein-labeled NAD^+^, providing direct evidence of active catalysis within the condensates (Fig. 5e,f). Time-course *in vitro* ADP-ribosylation assays demonstrated enhanced enzymatic activity following phase separation (Fig. 5g-i), indicating that condensate formation potentiates PARP10 activity beyond its basal level. To evaluate the consequences of condensation on enzymatic activity in cellular levels, we assessed auto-mono-ADP-ribosylation in condensation-deficient PARP10 mutants (3RE and 3EA). Both mutants exhibited significantly reduced auto-modification (Fig. 5j,k), indicating that phase separation is essential for optimal catalytic activity in cells.

Collectively, these findings demonstrate that PARP10 forms cytoplasmic condensates that are catalytically active both in cellular and reconstituted systems, and that phase separation enhances its ADP-ribosylation activity.

### PARP10 auto-ADP-ribosylation activity negatively regulates its phase separation

We wondered whether enzymatic activity could, in turn, regulate the formation and dynamics of PARP10 condensates. To investigate the role of enzymatic activity in PARP10 condensation, we generated a deletion mutant lacking the C-terminal catalytic domain (ΔCD). This mutant retained the ability to form cytoplasmic condensates in cells (Fig. 6a). Similarly, a point mutation (G888W) adjacent to the catalytic triad^36^, which abolishes PARP10 catalytic activity, did not impair condensation (Fig. 6a), indicating that ADP-ribosylation activity is dispensable for PARP10 phase separation. Using both ΔCD and G888W as catalytically inactive models, we compared their condensation properties with those of WT PARP10 at comparable expression levels (Fig. 6b). Notably, catalytic dead mutants formed larger condensates than WT (Fig. 6a). The partition coefficient (PC) of these mutants was significantly higher than that of WT (Fig. 6c), and their condensation occurred at a lower critical concentration (Fig. 6d), suggesting enhanced phase separation propensity. FRAP assays revealed reduced dynamics in condensates formed by the G888W and another catalytic deficient mutant H887E compared to WT (Fig. 6e,f). These findings imply that loss of catalytic activity enhances PARP10 condensation capacity, suggesting that ADP-ribosylation acts as an intrinsic suppressor of phase separation.

**Fig. 6.**
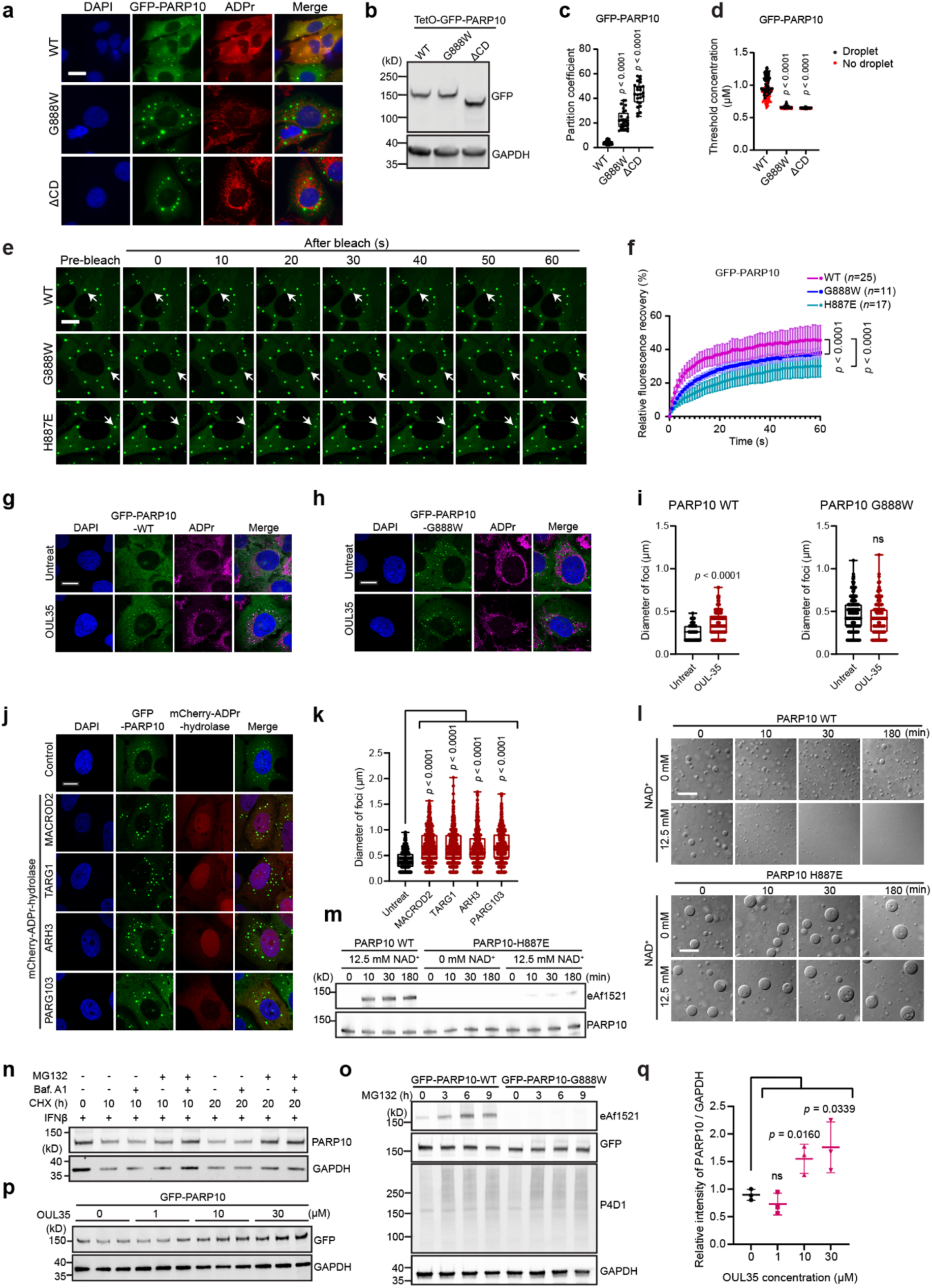
PARP10 condensates are regulated by ADP-ribosylation. **a**, Representative images of cells stably expressing GFP-PARP10 WT and catalytic deficient mutants (G888W and ΔCD). ADPr signals stained by eAf1521. **b**, Western blot showing the relative protein expression of each construct used in (**a**). **c**, Box plot for the partition coefficient (PC) of PARP10 WT and mutants in (**a**). Each symbol represents one condensate. At least 25 condensates were measured and analyzed. **d**, Phase separation threshold concentration was measured for PARP10 WT and mutants in (**a**). The absolute concentration in cells is converted by a standard curve created by measuring the intensity of recombinant PARP10 protein. Each symbol represents one cell. At least 100 cells were measured and analyzed. Each black dot indicates one cell with condensates, the red dot indicates the cell without condensates, and the line marks the threshold concentration for condensate formation. **e**,**f**, Representative images (**e**) and quantification (**f**) for FRAP experiments on PARP10 WT (*n* = 25), G888W (*n* = 11), and H887E (*n* = 17). The arrow indicated the bleached foci. Data are shown as mean ± SD. **g**,**h**, Representative images of GFP-PARP10 WT (**g**) and G888W (**h**) mutant treated with PARP10 inhibitor OUL-35. **i**, Box plot quantification of the diameter of PARP10 condensate in (**g**) and (**h**). **j**, Representative images of GFP-PARP10 co-expression with mCherry-MACROD2/TARG1/ARH3/PARG103. **k**, Box plot quantification of the diameter of PARP10 condensate in (**j**). Data are shown as mean ± SD. **l**, Representative images of PARP10 WT/H887E condensation induced by 10% Ficoll with or without NAD^+^ addition. **m**, Western blot showing the ADP-ribosylation and protein levels of PARP10 WT and H887E in (**l**). **n**, Western blot showing CHX chase assay for IFNβ-induced endogenous PARP10 protein level in U2OS cells coupled with MG132 or Bafilomycin A1 treatments. **o**, Western blot showing ADP-ribosylation of GFP-PARP10 WT in cells coupled with MG132 treatment for the indicated times. The accumulation of ubiquitin signal was detected with P4D1 ubiquitin antibody. The catalytic deficient mutant GFP-PARP10 G888W serves as a negative control. **p**, Western blot showing the accumulation of GFP-PARP10 protein in cells treated with different concentrations of PARP10 activity inhibitor OUL35. **q**, Quantification of the GFP-PARP10 protein levels in (**p**). Data are shown as mean ± SD. in (**q**). Scale bar: 10 μm for (**a**), (**e**), (**g**), (**h**), (**j**). 20 μm for (**l**).

To further examine the role of auto-ADP-ribosylation on PARP10 condensation, we treated cells with PARP10 inhibitor OUL-35 and found increased number and size of WT PARP10 condensates but without effect on the G888W condensates (Fig. 6g-i), supporting a specific role for catalytic activity in modulating condensation.

Given the reversibility of ADP-ribosylation, we assessed the impact of ADPr hydrolases on PARP10 condensation, including TARG1, MARCOD2, PARG103, and ARH3 (Extended Data Fig. 5a,b). Overexpression of each hydrolase in WT PARP10 stable cells increased condensate size (Fig. 6j,k), consistent with the removal of auto-ADP-ribosylation, which promotes phase separation. The negative regulatory mechanism was recapitulated *in vitro*, where Ficoll-induced PARP10 condensation was reversed in a time-dependent manner by auto-ADP-ribosylation induced by the addition of NAD^+^ (Fig. 6l). Western blot analysis confirmed PARP10 ADPr modification under these conditions (Fig. 6m). Importantly, NAD^+^ addition failed to dissolve condensates formed by catalytic dead mutants (Fig. 6l,m), confirming that the condensation suppression effect depends on the intrinsic enzymatic activity of PARP10.

Together, these results demonstrate that PARP10 possesses an intrinsic condensation capacity and reveal a negative feedback loop in which auto-ADP-ribosylation limits its own phase separation, both in cellular and reconstituted systems.

### Modified PARP10 is preferentially degraded via the ubiquitin-proteasome pathway

We next sought to elucidate the biochemical mechanism underlying the negative feedback loop in PARP10 condensation mediated by ADP-ribosylation. Our screening of PARP10 domains revealed a crucial role for ubiquitin-interacting motifs (UIMs) in PARP10 condensation (Fig. 3c,d). Polyubiquitin chains were enriched within PARP10 condensates, as detected by FK2 antibody staining (Extended Data Fig. 5c). These findings led us to propose that ADP-ribosylation might be closely associated with protein stability and regulates condensate dynamics. To investigate the involvement of ubiquitin signaling in PARP10 turnover, we assessed whether PARP10 is degraded through ubiquitin-dependent pathways. CHX chase assays were performed with MG132 or Bafilomycin A1 to inhibit the ubiquitin-proteasome system (UPS) or autophagy, respectively. PARP10 clearance was blocked by MG132 but not by Bafilomycin A1 (Fig. 6n), indicating that PARP10 degradation is mediated primarily by the UPS rather than autophagy. Using tandem ubiquitin-binding entities (TUBE) enrichment assays^37,38^, we detected ubiquitination of endogenous PARP10 following IFNβ stimulation and MG132-mediated inhibition of proteasomal degradation (Extended Data Fig. 5d). Similarly, exogenously expressed GFP-PARP10 was also ubiquitinated in cells (Extended Data Fig. 5e). Notably, short-term MG132 treatment resulted in a marked accumulation of ADP-ribosylated PARP10, while total PARP10 protein levels remained unchanged (Fig. 6o), suggesting that ADPr-modified PARP10 is selectively targeted for degradation. Consistent with this, treatment with the PARP10 inhibitor OUL35 led to a dose-dependent increase in PARP10 protein levels (Fig. 6p,q). Furthermore, WT PARP10 condensates dissolved upon cycloheximide (CHX) treatment, whereas G888W mutant condensates remained stable (Extended Data Fig. 5f,g). The increased solubility of ADPr-modified PARP10 compared to its unmodified form correlates with its accelerated turnover via the UPS. These findings support our earlier observation that ADP-ribosylation reduces PARP10’s capacity for condensation, reinforcing the hypothesis that enhanced auto-ADP-ribosylation acts as a negative feedback mechanism to disassemble PARP10 condensates.

To identify potential E3 ubiquitin ligases responsible for PARP10 ubiquitination, we examined members of the DTX family, known to interact with several PARP proteins^39–41^. Colocalization analyses using fluorescent reporters for DTX2, DTX3, DTX3L, and DTX4, as well as immunostaining for DTX1, revealed that all DTX family members colocalized with PARP10 condensates (Extended Data Fig. 6a-e). However, colocalization with DTX1 and DTX2 was abolished in the G888W mutant, suggesting an ADPr-dependent interaction, whereas DTX3, DTX3L, and DTX4 still associated with mutant condensates. Given that the WWE and DTC domains of DTX2 are implicated in ADPr recognition^42^, we generated a DTX2 mutant with five substitutions in these regions that disrupt ADPr binding. This mutant failed to colocalize with PARP10 condensates and exhibited markedly reduced interaction with PARP10 compared to WT DTX2 (Extended Data Fig. 6f,g), further supporting an ADPr-dependent recruitment mechanism. Additionally, our recent proteomic studies identified several TRIM E3 ligases^43^, including TRIM6 and TRIM38, as strong interactors of PARP10. We confirmed the colocalization of PARP10 condensates with TRIM6 and TRIM38 in cells (Extended Data Fig. 6h). Moreover, the association between PARP10 condensates and TRIM proteins was highly dependent on the presence of polyubiquitin chains (Extended Data Fig. 6h).

Collectively, these results demonstrate a close and multifaceted relationship between PARP10 auto-ADP-ribosylation and ubiquitination. The accumulation of ADPr modifications, combined with intrinsic ubiquitin-binding properties within the condensates, likely serves as a signal for the recruitment of multiple E3 ubiquitin ligases, thereby promoting the selective degradation of ADPr-modified PARP10.

### Condensation and enzymatic activity of PARP10 are essential for its antiviral function

To investigate whether enzymatic activity and condensation features of PARP10 are required for its antiviral function, we first employed the HEK293T cell model. Expression of WT PARP10 significantly inhibited VSV replication (Fig. 7a). In contrast, the catalytic dead mutant G888W and the condensation-deficient 3RE mutant exhibited markedly reduced antiviral activity (Fig. 7a), indicating that both the enzymatic activity and the capacity for condensation are necessary for PARP10-mediated suppression of VSV replication.

**Fig. 7.**
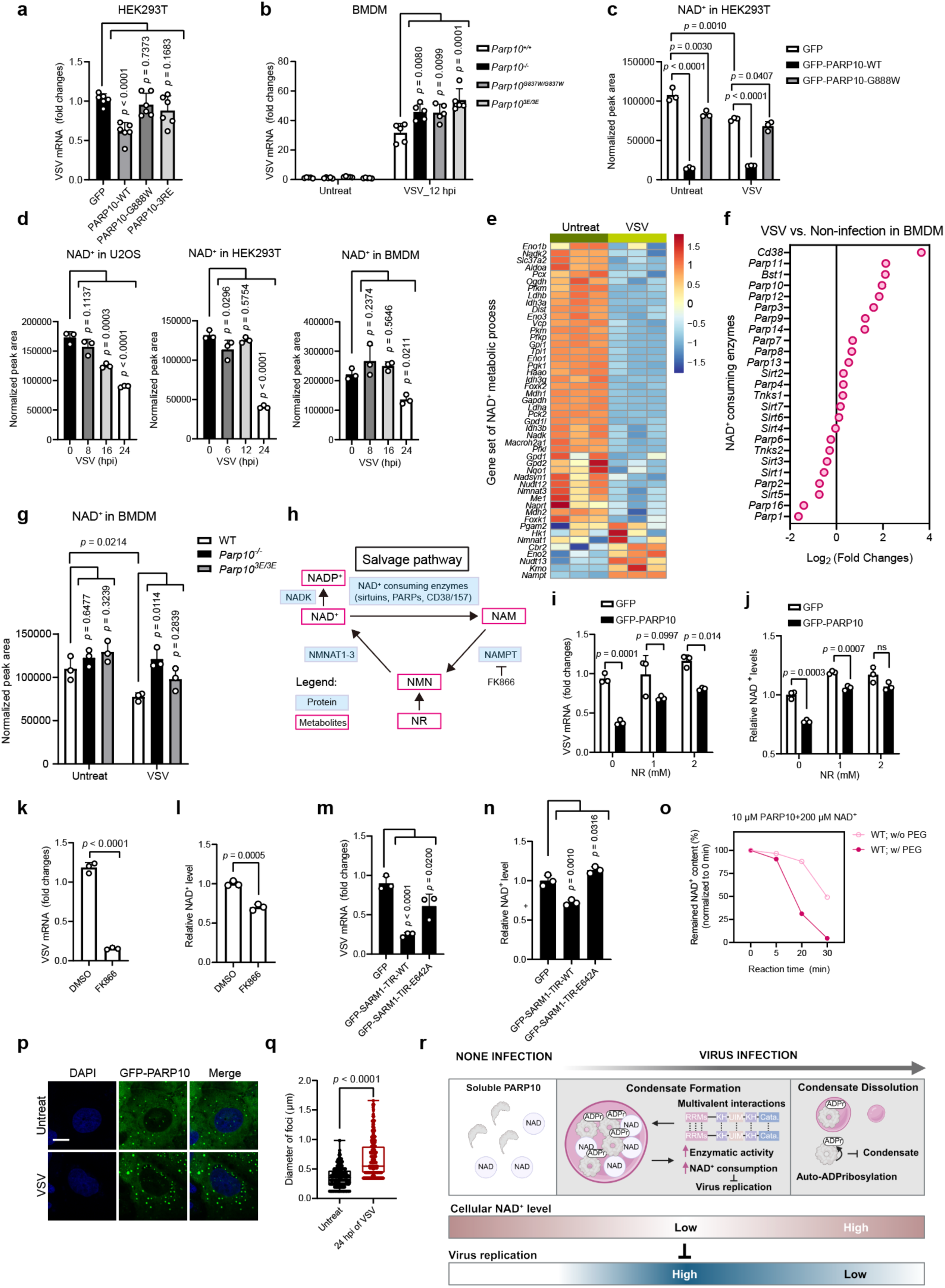
PARP10 condensates restrict VSV infection via regulating NAD^+^ homeostasis. **a**, Normalized VSV mRNA levels in HEK293T cells transfected with GFP/GFP-PARP10 WT/G888W/3RE and infected with VSV (8 hpi, MOI = 1.5). *n* = 6. **b**, Normalized VSV mRNA levels in *Parp10^+/+^*, *Parp10^-/-^*, *Parp10^G837W/G837W^* and *Parp10^3E/3E^* BMDM with or without VSV infection (12 hpi, MOI = 1.5). *n* = 5. **c**, NAD^+^ levels in HEK293T cells transfected with GFP/GFP-PARP10 WT/G888W, with or without VSV infection (8 hpi, MOI = 1.5), measured by MS. *n* = 3. **d**, NAD^+^ levels in WT U2OS, HEK293T and BMDM cells infected with VSV (MOI = 1.5) for the indicated times, measured by MS. *n* = 3. **e**, Heatmap of gene expression in BMDM infected with VSV (12 hpi, MOI = 1.5) or not. *n* = 3. Genes related to NAD^+^ metabolic process were shown. **f**, Fold changes of NAD^+^ consuming genes in BMDM with VSV (12 hpi, MOI = 1.5) infection relative to non-infected control. *n* = 3. **g**, NAD^+^ levels in *Parp10^+/+^* (WT), *Parp10^-/-^*, and *Parp10^3E/3E^* BMDM with or without VSV infection (12 hpi, MOI = 1.5), measured by MS. *n* = 3. **h**, Schematic diagram of the NAD^+^ salvage pathway and the effect of NAMPT inhibitor FK866. **i**, Normalized VSV mRNA levels in HEK293T cells transfected with GFP/GFP-PARP10 WT, followed by the supplement of the indicated concentrations of NR and VSV infection (8 hpi, MOI = 1.5). *n* = 3. **j**, NAD^+^ levels of cells in (**i**), determined by bioluminescent assays. *n* = 3. **k**, Normalized VSV mRNA levels in HEK293T cells treated with FK866 or DMSO (control), followed by VSV infection (8 hpi, MOI = 1.5). *n* = 3. **l**, NAD^+^ levels of cells in (**k**), determined by bioluminescent assays. *n* = 3. **m**, Normalized VSV mRNA levels in HEK293T cells transfected with GFP vector/GFP-SARM1-TIR WT/E642A, followed by VSV infection (8 hpi, MOI = 1.5). *n* = 3. **n**, NAD^+^ levels of cells in (**m**), determined by bioluminescent assays. *n* = 3. **o**, NAD^+^ consumption by recombinant PARP10 with or without the addition of PEG-8000 for condensation induction *in vitro* over time, determined by bioluminescent assays. **p**, Representative images of GFP-PARP10 condensate with or without VSV infection (24 hpi, MOI = 1.5). Scale bar: 10 μm. **q**, Box plot quantification of the diameter of PARP10 condensate in (**p**). **r**, Schematic diagram showing that PARP10 inhibits VSV infection by regulating NAD^+^ homeostasis. Data are shown as mean ± SD. in (**a**)-(**d**), (**g**), (**i**)-(**n**).

To examine the conservation of these features in the mouse ortholog, we generated a GFP-tagged mouse PARP10 (Mm_PARP10) reporter and expressed it in U2OS cells. Mm_PARP10 formed condensates similar to its human counterpart (Extended Data Fig. 7a). The G837W mutation in Mm_PARP10, equivalent to the G888W mutation in human PARP10, also enhanced condensation (Extended Data Fig. 7a). The 3R motif in the N-terminal RNA recognition motif (RRM) domain of human PARP10 corresponds to the HRR residues in Mm_PARP10, and the ubiquitin-interacting motif (UIM) is conserved between species. Consistent with the human protein, substitution of HRR to 3E or deletion of the UIM domain abolished Mm_PARP10 condensation (Extended Data Fig. 7a), indicating that both the functional properties and condensation behavior of PARP10 are evolutionarily conserved.

To further validate these findings *in vivo*, we generated two knock-in (KI) mouse models: *Parp10^3E/3E^* mice, which express a condensation-deficient form of PARP10, and *Parp10^G837W/G837W^* mice, which carry a catalytically inactive mutation (Extended Data Fig. 7b). The G837W mutation resulted in reduced PARP10 protein stability without affecting mRNA levels (Extended Data Fig. 7c,d). VSV replication was increased in BMDMs isolated from *Parp10^3E/3E^* KI mice compared to WT BMDMs (Fig. 7b).

Collectively, data from both cell-based assays and genetically modified mouse models demonstrate that the antiviral function of PARP10 depends on its ability to undergo phase separation and its catalytic activity.

### PARP10 exerts antiviral effects by modulating NAD^+^ homeostasis

Virus infection profoundly remodels the host cell metabolism, including alterations in NAD^+^ levels^44,45^. Previous studies have demonstrated that NAD^+^ levels decline following viral infection, including infection with SARS-CoV-2^46–48^. Given this metabolic perturbation in immune regulation, we investigated whether the antiviral activity of PARP10 is mediated through regulation of NAD^+^ metabolism. We first assessed NAD^+^ levels in HEK293T cells expressing PARP10 and observed a ∼7-fold reduction in NAD^+^ upon PARP10 expression (Fig. 7c). This decrease was abolished in cells expressing the catalytically inactive PARP10 G888W mutant (Fig. 7c), indicating that NAD⁺ consumption depends on PARP10 enzymatic activity. VSV infection alone also reduced NAD⁺ levels, although to a lesser extent than PARP10 overexpression (Fig. 7c).

To further explore the metabolic impact of PARP10, we conducted a metabolomic analysis in HEK293T cells transfected with WT PARP10, the catalytic dead mutant G888W, or the condensation-deficient 3RE mutant, with or without VSV infection. WT PARP10 expression induced extensive metabolic reprogramming, characterized by significant reductions in amino acid biosynthesis, TCA cycle intermediates, and taurine metabolism, alongside marked increases in amino-sugar, nucleotide-sugar, fructose, and mannose metabolism (Extended Data Fig. 7e,f). These metabolic changes were largely abrogated in the G888W mutant, resembling the GFP control, indicating dependence on catalytic activity. The 3RE mutant exhibited an intermediate phenotype, suggesting that PARP10 condensation also contributes to metabolic remodeling. Notably, VSV infection recapitulated many of the metabolic alterations observed with PARP10 expression (Fig. 7f), supporting a role for PARP10 in infection-induced metabolic reprogramming.

To assess the role of NAD^+^ metabolism in PARP10-mediated antiviral responses, we measured NAD^+^ levels at various time points after post-VSV infection in U2OS, HEK293T, and BMDMs. NAD^+^ level decreased consistently across all three cell types (Fig. 7d). RNA sequencing of BMDMs before and after VSV infection revealed dysregulation of genes involved in NAD^+^ biosynthesis and metabolism (Fig. 7e and Supplementary Table 3). Notably, NAMPT, a key enzyme in the NAD⁺ salvage pathway, was upregulated, suggesting a compensatory response to NAD⁺ depletion and cellular energy stress. This observation aligns with recent evidence positioning NAMPT as a metabolic sensor linking NAD⁺ dynamics to cellular energy status^49^. Major NAD^+^-consuming enzymes include cellular PARPs, Sirtuins, and CD38/CD157 ectoenzymes^50^. Our RNA-seq analysis showed upregulation of several NAD^+^-consuming enzymes during VSV infection, including *Cd38*, *Cd157(Bst1), Parp10*, and *Parp11* (Fig. 7f). Given the low expression of *Cd38* and *Cd157* in macrophages, their extracellular NAD^+^ hydrolase activity^51–53^, and the nuclear localization of endogenous PARP11, we hypothesized that cytoplasmic PARP10 is a primary contributor to NAD^+^ consumption during infection. To test this, we measured NAD^+^ levels in WT, *Parp10* KO, and *Papr10^3E/3E^*KI BMDMs following VSV infection. NAD^+^ depletion was significantly attenuated in *Parp10* KO cells, and less effectively restored in *Parp10^3E/3E^* mutant (Fig. 7g), indicating that PARP10 activity and, to a lesser extent, the condensation, are essential for infection-induced NAD^+^ decline.

NAD⁺ depletion has been shown to function as an antiviral mechanism in bacterial-phage systems^17^. We hypothesized that a similar strategy may be conserved in mammalian antiviral immunity. To evaluate the functional significance of NAD⁺ consumption, we manipulated NAD⁺ levels using the cell-permeable precursor nicotinamide riboside (NR) to boost NAD⁺ and the NAMPT inhibitor FK866 to suppress NAD⁺ synthesis (Fig. 7h). NR treatment dose-dependently reversed the antiviral effect of PARP10 (Fig. 7i,j), while FK866-mediated NAD⁺ reduction strongly inhibited viral replication (Fig. 7k,l). Consistent with this, expression of the TIR domain of SARM1^24,54^, a well-characterized NAD⁺-cleaving enzyme in mammals, led to NAD⁺ depletion (Fig. 7n) and robust suppression of VSV replication (Fig. 7m). This antiviral effect was dependent on catalytic activity, as the SARM1 E642A mutant failed to reduce NAD⁺ levels or inhibit viral replication (Fig. 7m,n).

We further examined the role of PARP10 condensation in NAD⁺ consumption using an *in vitro* assay. PEG-induced phase separation of PARP10 markedly accelerated NAD⁺ consumption kinetics (Fig. 7o), correlating with enhanced auto-ADP-ribosylation (Fig. 5i). During late-stage VSV infection (24 hours post-infection), PARP10 auto-ADP-ribosylation decreased (Extended Data Fig. 7g), likely due to NAD⁺ exhaustion, while PARP10 condensates increased in size (Fig. 7p,q), suggesting a feedback mechanism linking substrate availability to condensate dynamics.

Gene Set Enrichment Analysis (GSEA) comparing transcriptomes of PARP10-expressing and control cells revealed significant enrichment of gene signatures upregulated during SARS-CoV-2 infection in ACE2-expressing A549 cells^55^ (Extended Data Fig. 7h,i and Supplementary Table 4). To extend these findings to human disease, we analyzed publicly available RNA-seq data from healthy individuals and patients with COVID-19^56^. Most genes involved in NAD⁺ catalytic processes were downregulated in COVID-19 patients, whereas key components of the NAD⁺ salvage pathway, such as NAMPT, NMNAT1, and NMNAT2, were upregulated, consistent with a compensatory response (Extended Data Fig. 7j). A similar trend was observed for NAD⁺-consuming enzymes (Extended Data Fig. 7k), mirroring our cellular model (Fig. 7j,k). Among the ARTD family, PARP10, PARP9, PARP12, PARP13 (ZC3HAV1), and PARP14 were markedly induced in COVID-19 patients, whereas PARP16 and members of Sirtuins remained stable or were slightly downregulated (Extended Data Fig. 7k,l).

Collectively, these findings demonstrate that the antiviral function of PARP10 is dependent on both its enzymatic activity and its capacity to undergo phase separation. By directly modulating NAD⁺ homeostasis, PARP10 plays a pivotal role in antiviral defense and has significant implications for understanding metabolic regulation in viral infections, including COVID-19.

## Discussion

Here, we revealed the importance of PARP10 condensation during virus infection using cell and mouse models. Characterization of PARP10 condensation revealed the biochemical basis for PARP10 condensation and suggested a potentially broader regulatory principle for the compartmentalization of other ARTD enzymes. The compartmentalization of ARTD enzymes through condensation provides a novel mechanism for regulating NAD^+^ metabolism, which is crucial for maintaining the cellular oxidative state, regulating energy metabolism, promoting cell proliferation, and modulating immune responses. Our work on PARP10 condensate, from cell biology, biochemistry, and structural biology perspectives, highlighted the intrinsic feature of higher-order assembly.

Redundancy among ARTD family members likely exists during virus-induced immune responses, as we observed concurrent upregulation of PARP10, PARP13, and PARP14. Prior studies have reported the recruitment of several ARTDs including TNKS1, PARP12, PARP13 and PARP15 to stress granules (SGs), cytoplasmic compartments critical in host–virus interactions^57,58^. Notably, our study identifies PARP10 as forming a distinct cytoplasmic condensate that is separate from SG-associated ARTDs.

Multivalent interactions mediated by multiple self-association domains are critical for PARP10 condensation. PARP10 contains several structured RRM and KH domains that collectively drive its condensation in cells. Cryo-EM structural analysis combined with mutagenesis experiments demonstrates that an arginine-rich tract is essential for PARP10 oligomerization and phase separation both *in vitro* and in cells. Recent structural studies on TNKS1 have revealed diverse conformations involving head-to-head or tail-to-tail interaction between catalytic domain, which can regulate the catalytic activity^59^. Further investigations will elucidate the precise inter-domain interactions, in addition to the arginine tract identified here, that govern m PARP10 oligomerization.

Furthermore, we show that PARP10 undergoes ubiquitination in cells, which regulates its protein stability. This auto-ubiquitination of PARP10 is supported by recent evidence indicating that a mono-ADP-ribosyl ubiquitylation (MARUbylation) mechanism contributes to its modification^41^. Our work uncovers a diverse and flexible interaction network between PARP10 and multiple E3 ubiquitin ligases, including members of the DTX and TRIM families. However, the specific E3 ligases responsible for PARP10 ubiquitination remain to be identified. We observe faster degradation kinetics of ADP-ribosylated PARP10 compared to its catalytically inactive mutant, mediated by the ubiquitin–proteasome system (UPS). Combined with our finding that auto-ADP-ribosylation promotes disassembly of PARP10 condensates, these results suggest a negative feedback loop regulating PARP10 condensation. Thus, PARP10 condensation and self-modification constitute a self-limiting mechanism that fine-tunes its interferon-stimulated antiviral activity during viral infection (Fig. 7r). This regulatory circuit may be evolutionarily conserved and applicable to other interferon-stimulated genes (ISGs) activated in response to viral invasion.

We demonstrate that PARP10 restricts viral replication through NAD^+^ consumption. NAD^+^ depletion has been established as an antiviral strategy in the bacterial-phage system^17^. In contrast to the near-complete NAD^+^ depletion in prokaryotes, PARP10 reduces mammalian NAD^+^ levels only by a few fold (Fig. 7c,g). Given that basal NAD^+^ concentrations in mammalian cells are approximately ∼100 μM across various cell types, such partial depletion is unlikely to directly mediate antiviral effects. Instead, a signaling pathway that senses NAD^+^ levels may underlie PARP10’s antiviral function. Recent evidence indicates that NAD^+^-sensing pathways can induce antiviral gene expression in plants^60^. While the ADP-ribosyltransferase activity of PARP10 is well characterized, the potential production of a novel signaling nucleotide by PARP10 remains unexplored. Notably, a recent study by Rotem Sorek’s group demonstrated that the SARM1 TIR domain generates cyclic ADPR as a minor product^61^, which contributes to SARM1-mediated axonal degeneration in mammals.

Although NAD^+^ consumption significantly contributes to the antiviral effects of PARP10, additional mechanisms may cooperate to enhance its antiviral function. One possibility is that PARP10 condensates may restrict viral assembly by sequestering and ADP-ribosylating the nucleocapsid of VSV. Recent studies have shown that PARP10 mono-ADP-ribosylates alphaviral non-structural protein 2 (nsP2), inhibiting nsP2 protease enzymatic activity^62^. Similarly, PARP7 and PARP12 can ADP-ribosylate alphaviral nsP3 and nsP4, with the viral Macro domain-containing nsP3 counteracting this antiviral effect^63^. Future studies will determine whether PARP10 directly binds and modifies viral capsid components, and how such modifications influence virion assembly.

Cellular NAD^+^ homeostasis is fundamental to nucleotide and energy metabolism. Enzymes from bacteria and plants can cleave NAD^+^ or convert ATP to ITP, thereby manipulating host metabolic pathways. In mammals, ARTDs not only deplete NAD^+^ through protein upregulation and condensation but also selectively modify both host and pathogen substrates. Thus, the substrate specificity of individual ARTD enzymes may account for their differential antiviral potency. In light of recent discoveries positioning NAD^+^ as a central node in bacterial–phage conflict^18,64,65^, our findings underscore PARP10 as key components of the mammalian innate immune defense that target NAD^+^ metabolism.

## Acknowledgements

We thank Dr. Pilong Li from Tsinghua University and Dr. Liang Tao from Westlake University for technical support. We thank the technical support from the Westlake Biomedical Research Core Facility. The work was supported by grants from the National Key Research and Development Project of China (2025YFC3409700 to P.Y.), the National Natural Science Foundation of China (32470733 and 32170696 to P.Y.; 32271264 to Z.S.), and by the “Pioneer” and “Leading Goose” R&D Program of Zhejiang (2024SSYS0030).

## Author contributions

X.C. performed cellular assays, *in vitro* phase separation assay, enzymatic activity assay, virus infection, and mouse model studies. Q.X. and Z.S. performed the cryo-EM study and analyzed the data. Y.L., Z.Z., Z.W., Y.Z., Q.C., L.T., and Z.Y. participated in the plasmid construction, protein purification, mouse BMDM isolation and culture, RT-qPCR, and microscopy studies. P.Y. and Z.S. supervised the study. P.Y., X.C., and Z.S. wrote the manuscript with input from all authors.

## Competing interests

The authors declare no conflict of interest.

---

**Extended Data Fig. 1.**
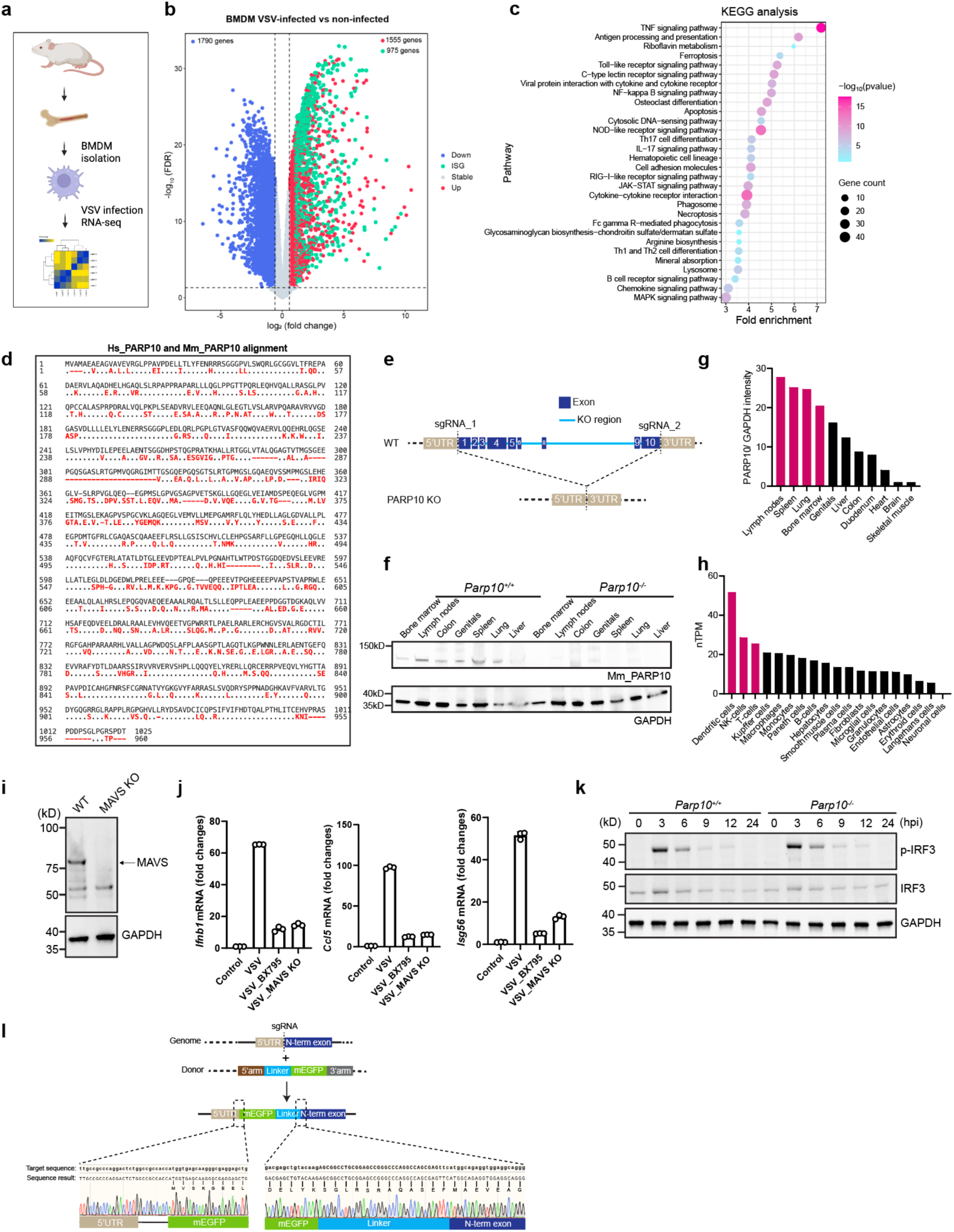
PARP10 is an antiviral factor. **a**, A flow chart for BMDM isolation and RNA-seq analysis, created through BioRender. **b**, Volcano plot showing the upregulated and downregulated genes in BMDM infected with VSV (12 hpi, MOI = 1.5) compared to non-infected control. Blue dot indicates downregulated genes, red dot indicates upregulated genes, green dot indicates upregulated interferon-stimulated genes (ISGs). **c**, KEGG analysis of upregulated genes in (**b**). **d**, Amino acid sequence comparison between human and mouse PARP10. **e**, Schematic diagram showing the KO of the mouse *Parp10* gene. **f**,**g**, Western blot showing the PARP10 protein expression in multiple tissues from *Parp10^+/+^* and *Parp10^-/-^*mice. Normalization to GAPDH is shown in (**g**). **h**, Relative expression of PARP10 from human single-cell RNA-seq data extracted from https://www.proteinatlas.org/. **i**, Western blot validating the MAVS KO in HEK293T cells. **j**, Normalized mRNA levels of *Ifnb1*/*Ccl5*/*Isg56* in WT/BX795-treated/MAVS KO HEK293T cells transfected with GFP vector or GFP-PARP10, followed by VSV infection (8 hpi, MOI = 1.5). *n* = 3. **k**, Western blot to determine the levels of p-IRF3, total IRF3, and PARP10 in *Parp10^+/+^* and *Parp10^-/-^* BMDM at different time points post-VSV (MOI = 1.5) infection. **l**, Schematic diagram showing the EGFP KI strategy for the generation of the *Parp10^mEGFP/mEGFP^* mice.

**Extended Data Fig. 2.**
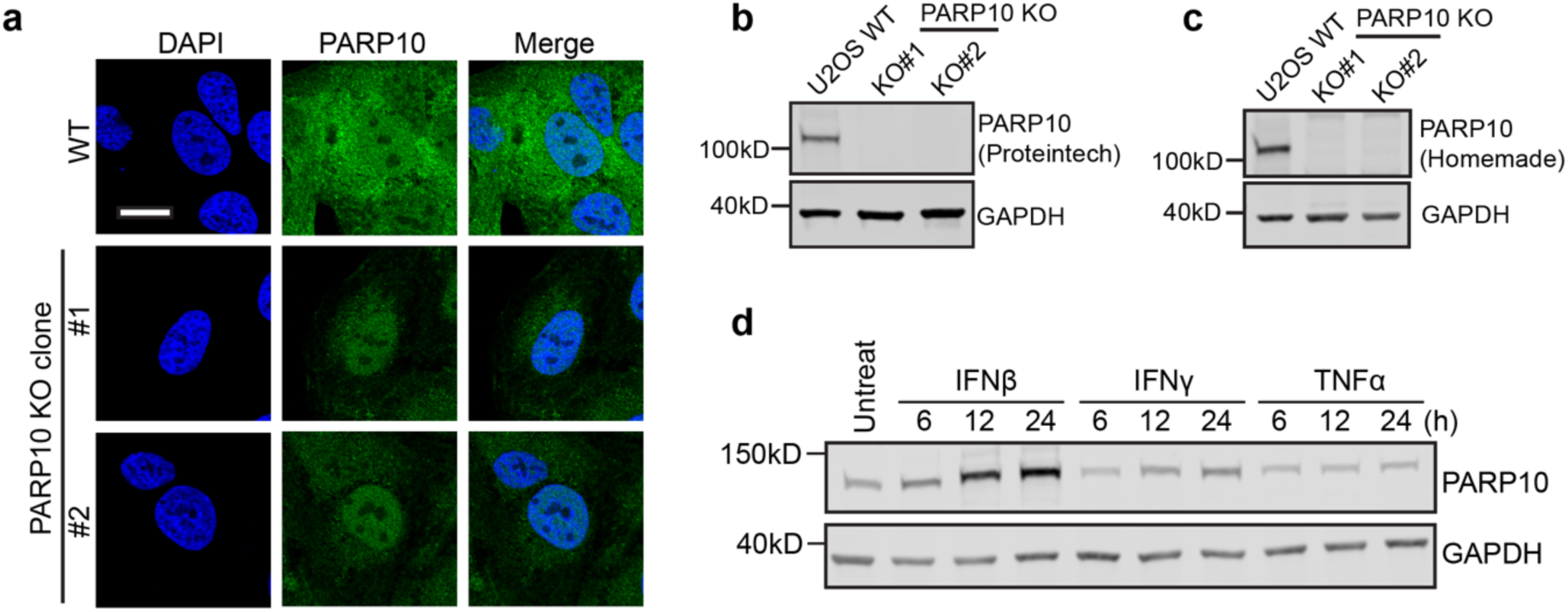
Generation of PARP10 knock-out cell line. **a**, Representative images of endogenous PARP10 staining in WT and PARP10 KO U2OS cells. **b**,**c**, Western blot showing the expression of endogenous PARP10 in U2OS cells and verification of PARP10 KO. PARP10 was stained with commercial antibody from Proteintech (**b**) and homemade antibody (**c**). **d**, Western blot showing the induction of endogenous PARP10 in U2OS with IFNβ, IFNγ, and TNFα stimulation. Scale bar: 10 μm for (**a**).

**Extended Data Fig. 3.**
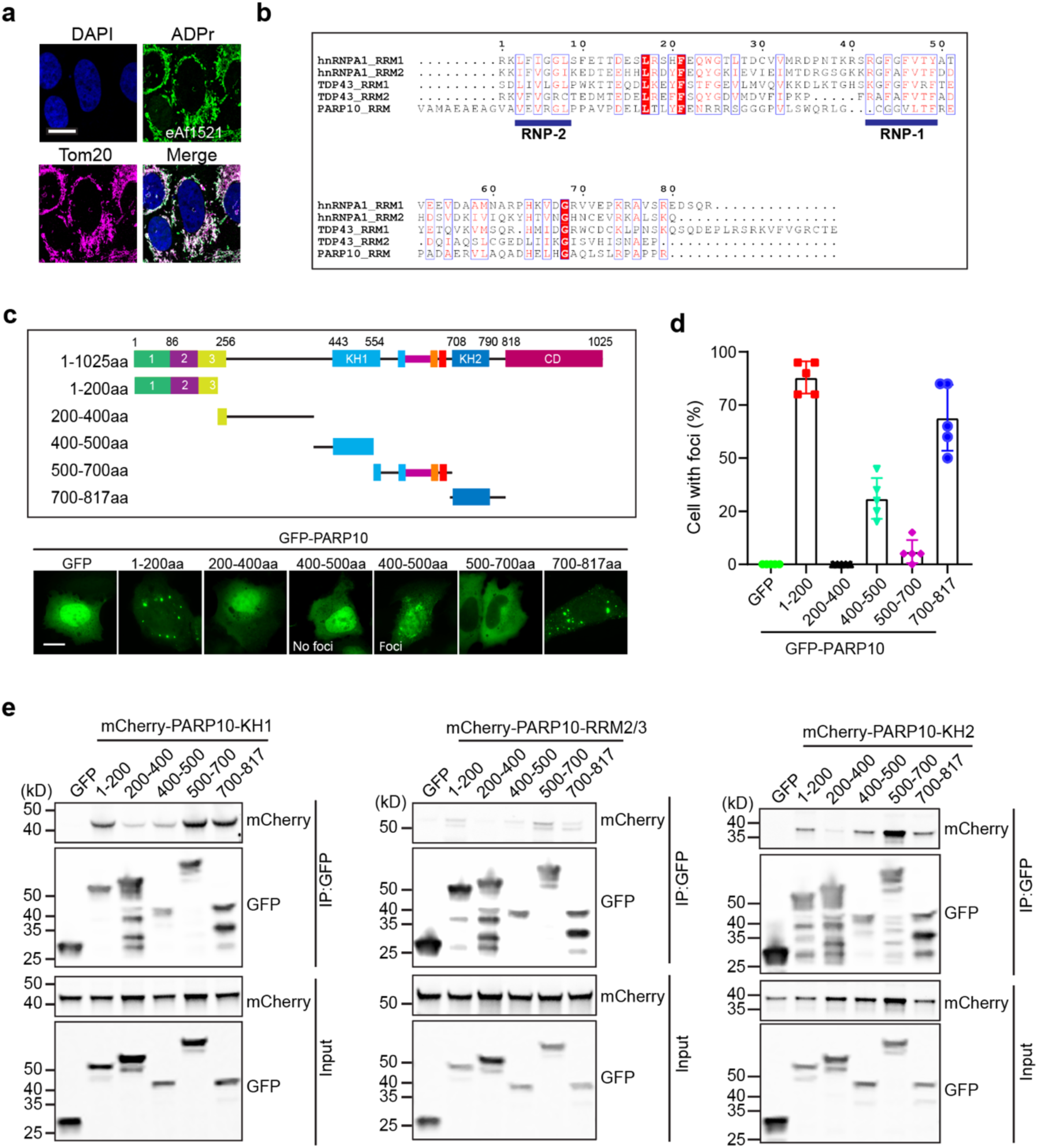
Multivalent interactions across PARP10. **a**, Representative images of ADPr staining with eAf1521 and mitochondria staining with Tom20. **b**, RRM sequence alignment of PARP10 with TDP43 and hnRNPA1. **c**, Diagram showing the constructs used in the assay, and the representative images are shown below for each construct. Scale bar: 10 μm. **d**, Quantification of percentage of cells with foci formation in (**c**). **e**, Western blot showing the interaction between PARP10 domains (RRM2/3, KH1, or KH2) and different fragments in (**c**), determined by co-immunoprecipitation assay.

**Extended Data Fig. 4.**
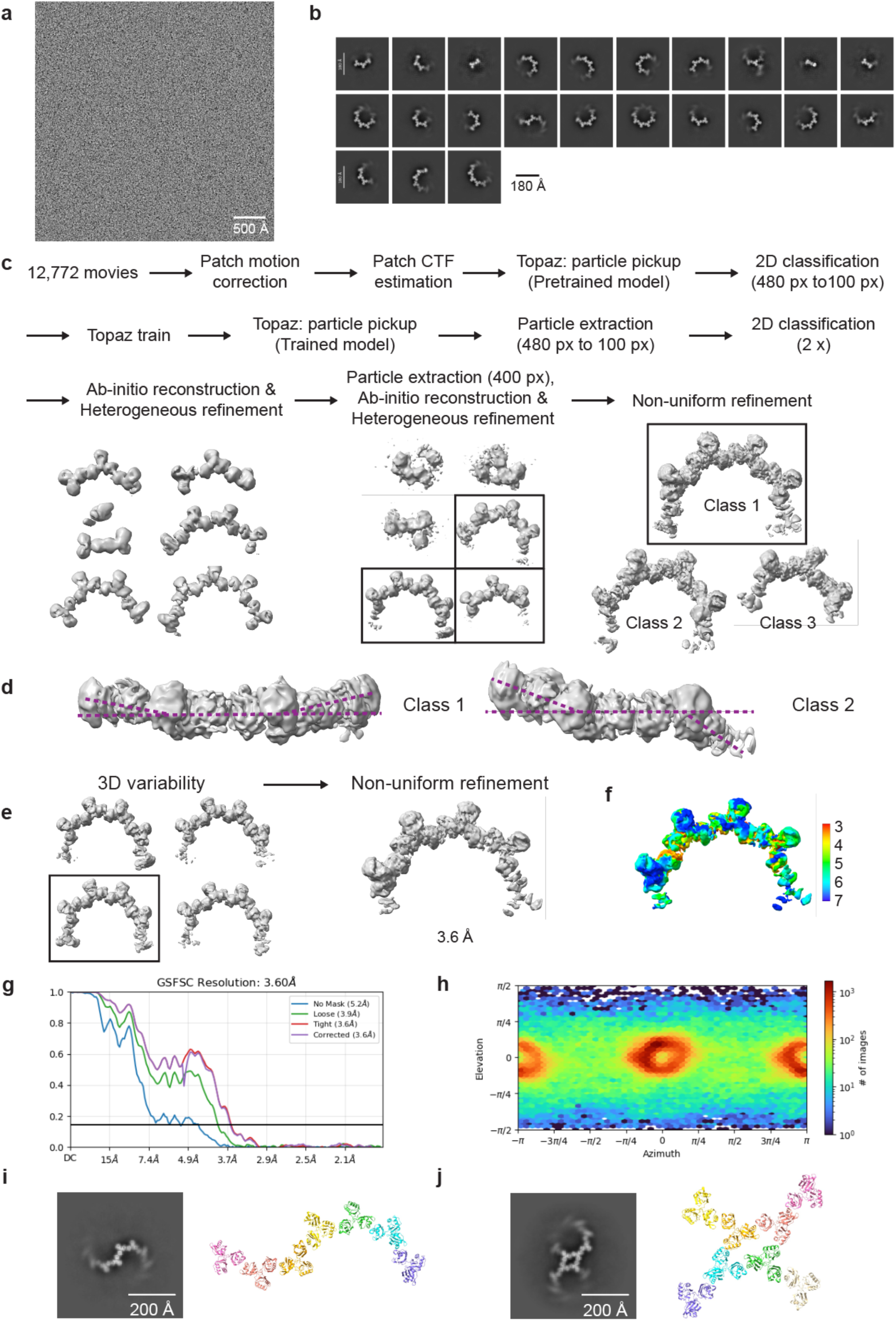
Cryo-EM analysis of PARP10 RRM domains. **a**, A representative cryo-EM micrograph of full-length PARP10 protein. Scale bar: 500 Å. **b**, 2D class averages were selected for 3D reconstruction. Scale bar: 180 Å. **c**, Flow chart for cryo-EM data analysis. **d**, Maps of two classes of PARP10 RRM rings with different twisted angles. **e**, 3D variability analysis and non-uniform refinement of RRM oligomers. **f**-**h**, Map local resolution (**f**), Fourier shell correlation (FSC) curves (**g**), and particle distribution used in 3D reconstruction (**h**) of PARP10 RRM1-3 pentamer. **i**,**j**, 2D classes and corresponding models to show the two types of higher-order oligomerization of RRM1-3. Scale bar: 200 Å.

**Extended Data Fig. 5.**
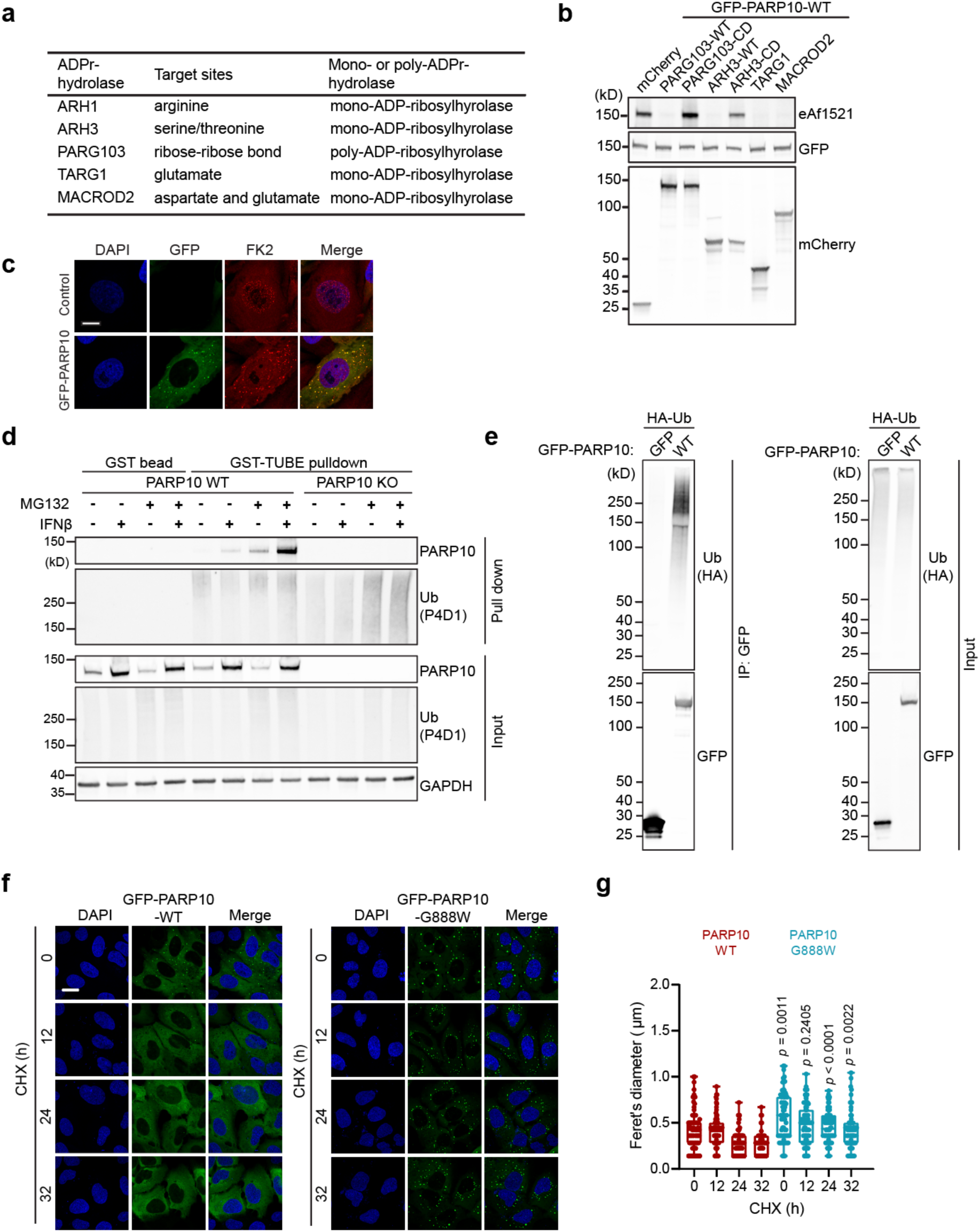
PARP10 condensates are regulated by ADP-ribosylation. **a**, List of ADPr hydrolases used in the study. **b**, Western blot showing the effect of mCherry-tagged ADPr hydrolase and its catalytic deficient (CD) mutants on the GFP-PARP10 auto-ADP-ribosylation in HEK293T cells. The auto-ADP-ribosylation was probed by eAf1521. **c**, Representative images of GFP-PARP10 condensates stained for FK2. **d**, TUBE pulldown assay showing the ubiquitination of PARP10 in response to MG132 and IFNβ treatment. GST bead only and PARP10 KO cells were used as controls. **e**, Western blot showing the ubiquitination of PARP10 in cells. **f**, Representative images of GFP-PARP10 WT/G888W stable cells treated with CHX for different times. **g**, Box plot quantification of the diameter of PARP10 condensate in (**f**). Scale bar: 10 μm for (**c**) and (**f**).

**Extended Data Fig. 6.**
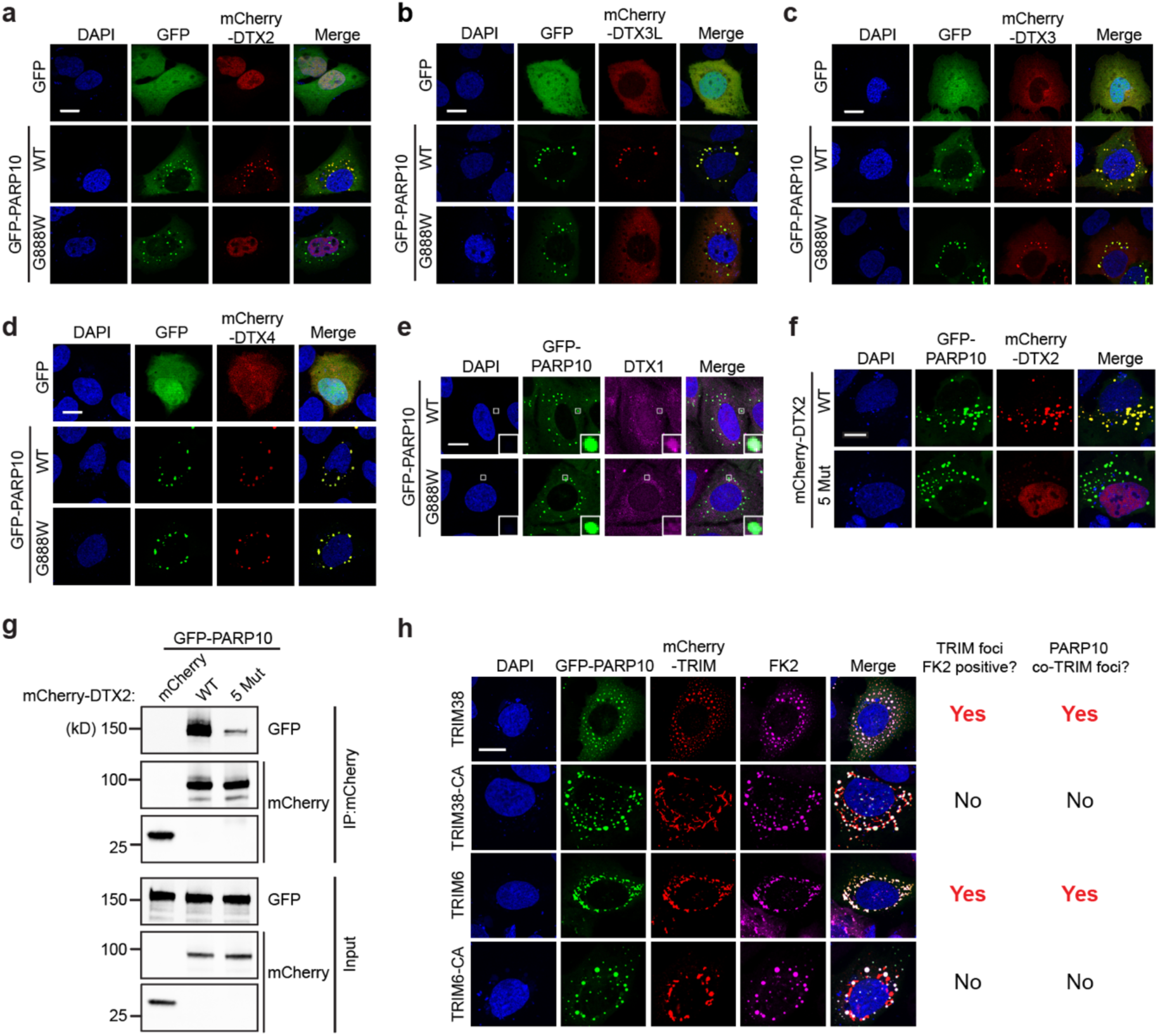
Diverse interaction with E3 ligases in PARP10 condensation. **a**-**d**, Representative images of cells co-transfected with GFP/GFP-PARP10 WT/G888W and mCherry-DTX2 (**a**), DTX3L (**b**), DTX3 (**c**), DTX4 (**d**). **e**, Representative images of cells transfected with GFP-PARP10 WT/G888W and stained with DTX1 antibody. **f**, Representative images of cells co-transfected with GFP-PARP10 WT and mCherry-DTX2 WT or 5 Mut which disrupts the ADPr binding. **g**, Western blot showing the interaction between GFP-PARP10 WT and mCherry/mCherry-DTX2 WT/5 Mut, determined by co-immunoprecipitation assay. **h**, Representative images and summary of GFP-PARP10 condensates colocalized with FK2-positive TRIMs, but not with the FK2-negative catalytic mutants. Scale bar: 10 μm for (**a**)-(**f**) and (**h**).

**Extended Data Fig. 7.**
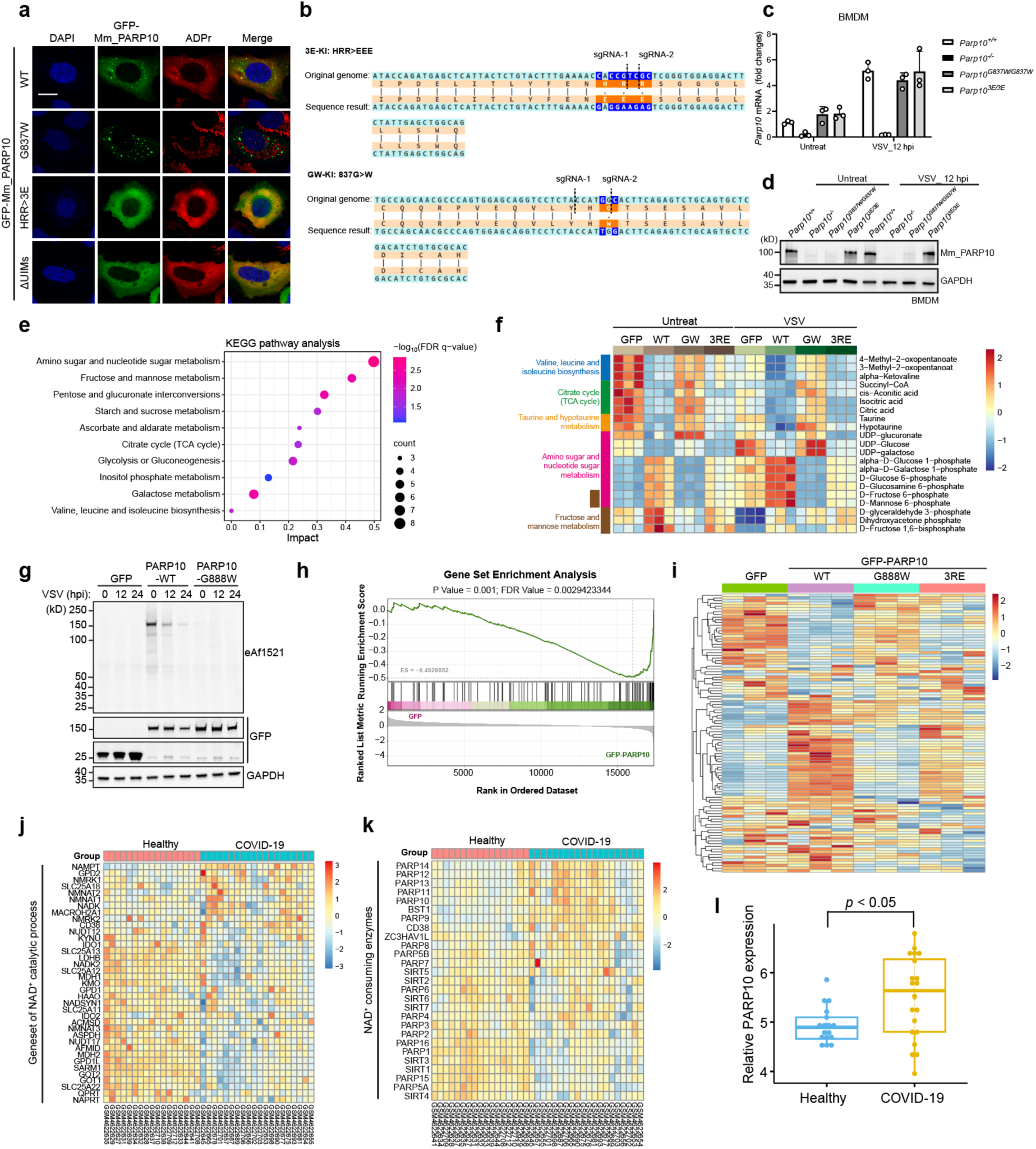
PARP10 condensates confer antiviral effects via regulating NAD^+^ homeostasis. **a**, Representative images of cells transfected with mouse PARP10 WT and mutants. ADPr signals were detected using eAf1521. Scale bar: 10 μm. **b**, Schematic diagram showing the generation of *Parp10^G837W/G837W^* and *Parp10^3E/3E^* mice with verification by Sanger sequencing. **c**, Normalized *Parp10* mRNA levels in *Parp10^+/+^*, *Parp10^-/-^*, *Parp10^G837W/G837W^*and *Parp10^3E/3E^* BMDM infected with VSV (12 hpi, MOI = 1.5) or not. *n* = 3. Data are shown as mean ± SD. **d**, Western blot showing the PARP10 protein expression in *Parp10^+/+^*, *Parp10^-/-^*, *Parp10^G837W/G837W^* and *Parp10^3E/3E^*BMDM infected with VSV (12 hpi, MOI = 1.5) or not. **e**, KEGG analysis of metabolism pathways affected by PARP10 expression in HEK293T cells. **f**, Heatmap of metabolite levels in HEK293T cells transfected with GFP/GFP-PARP10 WT/G888W/3RE, with or without VSV infection (8 hpi, MOI = 1.5), measured by MS. *n* = 3. Metabolites of particular interest are colored by category. Metabolites with significant changes (*p* < 0.05) are shown. **g**, Western blot showing the ADP-ribosylation of GFP-PARP10 in cells infected with VSV (MOI = 1.5) for the indicated times. Cells transfected with GFP vector and catalytic-deficient mutant (G888W) served as negative controls. PARP10 ADP-ribosylation was probed with eAf1521. **h**, GSEA showing the enriched gene set^55^ in HEK293T cells expressing GFP-PARP10 relative to GFP vector. **i**, Heat map of gene expression in (**h**) gene set. The expression profile for GFP, GFP-PARP10 WT, G888W, and 3RE mutants was shown. **j**,**k**, Heat map of gene expression related to NAD^+^ catalytic process (**j**) and NAD^+^ consumption (**k**) from the whole blood samples in healthy (*n* = 17) and COVID-19 (*n* = 20) populations.^56^ **l**, Comparison of PARP10 expression between healthy (*n* = 17) and COVID-19 (*n* = 20) populations. Data are shown as mean ± SD.

## Methods

### Antibodies

The following primary antibodies were used: Rabbit Polyclonal Anti-PARP10 (Proteintech, 26072-1-AP), Rabbit Polyclonal Anti-PARP2 (Proteintech, 26072-1-AP), Mouse Monoclonal Anti-PARP4 (Santa Cruz Biotechnology, sc-515898), Rabbit Monoclonal Anti-Poly/Mono-ADP-Ribose (E6F6A) (Cell Signaling Technology, 83732S), Mouse Monoclonal Anti-Poly-ADP-Ribose (10H) (Abcam, ab14459), Mouse Monoclonal Anti-GFP (Proteintech, 66002-1-Ig), Rabbit Polyclonal Anti-GFP (Proteintech, 50430-2-AP), Rabbit Polyclonal Anti-mCherry (Proteintech, 26765-1-AP), Mouse Monoclonal Anti-GAPDH (Proteintech, 60004-1-Ig), Mouse Monoclonal Anti-G3BP1 (BD Biosciences, 611126), Rabbit Polyclonal Anti-LSM14A (Proteintech, 18336-1-AP), Rabbit Polyclonal Anti-RAB7A (Proteintech, 55469-1-AP), Mouse Monoclonal Anti-TOM20 (Santa Cruz Biotechnology, sc-17764), Rabbit Polyclonal Anti-DTX1 (Proteintech, 18350-1-AP), Rabbit Polyclonal Anti-MAVS (Proteintech, 14341-1-AP), Mouse Monoclonal Anti-HA-tag (MBL International, M180-3), Mouse Monoclonal Anti-Ubiquitinylated Proteins (Clone FK2) (Millipore, 04–263), Mouse Monoclonal Anti-Ubiquitin (P4D1) (Cell Signaling Technology, 3936), Rabbit Monoclonal Anti-IRF3 (D83B9) (Cell Signaling Technology, 4302), Rabbit Monoclonal Anti-Phospho-IRF3 (Ser396) (4D4G) (Cell Signaling Technology, 4947). The secondary antibodies used in this study include Alexa Fluor 488 (A11029, A11008)/555 (A21428, A21422)/647 (A21235, A21245) secondary antibodies (Invitrogen) for immunofluorescence; IRDye 680RD (926-68070) and IRDye 800CW (926-32211) secondary antibodies (LI-COR) for immunoblot.

### Cell culture

U2OS, HEK293T, BHK-21, and BRS-T7/5 cells were cultured in Dulbecco’s modified Eagle’s medium (Thermo Fisher Scientific, 11995500CP) supplemented with 10% fetal bovine serum (ExCellBio, FSP500) and 1% penicillin/streptomycin (MACKLIN, Q6532). Bone-marrow-derived macrophages (BMDMs) were isolated from the tibia and femur of C57BL/6 mice. BMDM cells were cultured in DMEM with 10% fetal bovine serum (ExCellBio, FSP500), 1% penicillin/streptomycin (MACKLIN, Q6532), and 30% L929 supernatant at 37 °C for 7 days. Cells were all maintained at 37 °C in a humidified incubator with 5% CO_2_. The Hieff Trans™ Liposomal Transfection Reagent (YEASEN, 40802ES01) or PEI MAX (Polysciences, 24765-100) is used for transient transfection according to the manufacturer’s instructions. All cells are regularly checked for mycoplasma and are without contamination.

DH5α competent cells (TsingKe, TSC-C14) were used for plasmid amplification. Chemically competent bacteria strain Rosetta (DE3) (TIANGEN, CB108-02) was used for protein production. *E. coli* cells were cultured in LB media containing appropriate antibiotics at 37 °C in a shaker incubator (200 rpm).

### Mouse models

All mice were on C57BL/6 background. *Parp10^-/-^*, *Parp10^3E/3E^*, *Parp10^G837W/G837W^* and mEGFP-*Parp10* knock-in (*Parp10^mEGFP/mEGFP^*) mice were generated via CRISPR/Cas9 mediated gene editing by Cyagen Bioscience Inc. Mice were maintained under specific-pathogen-free conditions with controlled temperature (18–23 °C), humidity (40–60%) and a 12-hour light/12-hour dark cycle. The infectious virus-included animal experiments were performed in the physical containment level 2 laboratory with approval from the Institutional Animal Care and Use Committee of Westlake University (AP#23-110-3-YPG-3).

The gRNA sequences used are listed as follows:

*Parp10^-/-^*gRNA_1 target sequence:

5’-GCGGGCAATGGTGATGAGCA-3’

*Parp10^-/-^*KO gRNA_2 target sequence:

5’-TCTAGTCATATCTTCGGCTC-3’

*Parp10^3E/3E^*gRNA_1 target sequence:

5’-GTCCTCCACCCGAGCGACGG-3’

*Parp10^3E/3E^*gRNA_2 target sequence:

5’-CTTTGAAAACCACCGTCGCT-3’

*Parp10^G837W/G837W^*gRNA_1 target sequence:

5’-ACTCTGAAGTGCCATGGTAG-3’

*Parp10^G837W/G837W^*gRNA_2 target sequence:

5’-CTGCAGACTCTGAAGTGCCA-3’

*Parp10^mEGFP/mEGFP^*gRNA_1 target sequence:

5’-TCTGCCATCAGAGTCCTGGG-3’

*Parp10^mEGFP/mEGFP^*gRNA_2 target sequence:

5’-CACCTCTGCCATCAGAGTCC-3’

### Virus stock generation

The Semliki Forest Virus (SFV) was generated from a DNA vector driven by a CMV promoter, a gift from Dr. Andres Merits from University of Tartu, Estonia. The Vesicular Stomatitis Virus (VSV) DNA plasmid, driven by a T7 promoter, was purchased from Miaoling Biology. The viral stock was filter-sterilized with a 0.45-μm filter and stored at -80 °C. The viral concentration was determined by plaque assay using BHK-21 cells. Briefly, monolayers of BHK-21 cells in a 24-well plate were incubated with a series of tenfold-diluted virus samples for 48 h. The cells were fixed with 10% formaldehyde (Sigma-Aldrich, F8775) for 1 h and stained with 0.5% crystal violet (Rhwan, R019732) solution for 15 min. The plaques were counted manually.

### Viral infection in mouse models

8-12 weeks old mice were infected with VSV (1x10^8^ PFU/mouse) diluted in PBS (Biosharp, BL302A) by intraperitoneal injection, with PBS-only as control. At 24 h post of infection, the spleen and liver were obtained from each mouse and flash-frozen in liquid nitrogen. Tissues were ground under liquid nitrogen, and RNA was isolated with TRIzol™ reagent (Thermo Fisher Scientific, 15596018) for gene expression analysis.

### Cell stimulation

Cells were treated with 500 μM sodium arsenite (Merck, 1.06277.1003) for 1 h, 20 ng/mL human IFNβ (Peprotech, 300-02BC), IFNγ (Peprotech, 300-02), TNFα (Peprotech, 300-01A) for 24 h, 10 μM Olaparib (MCE, HY-10162) or OUL-35 (R&D, 6344) for 5 h, VSV at MOI=1.5 for indicated times, SFV at MOI=0.5 for indicated times, 50 μg/mL poly(I:C) LMW (Invivogen, tlrl-picw) for 5 h, 100 μg/mL RNAse A (Takara, 740505) in PBS, 0.5 μM LysoTracker (Thermo Fisher Scientific, L12492) for 30 min at 37 °C, 1 μM CellTracker CM-Dil Dye (Invitrogen, C7000) for 30 min at 37 °C, 10 μM BX795 (MCE, HY-10514) 3 h before and during viral infection, 1-2 mM Nicotinamide riboside chloride (NR) (MCE, HY-123033A) or 1 μM FK866 (MCE, HY-50876) 24 h before and during viral infection.

### Plasmid construction

DNA fragments encoding human PARP10 were amplified from U2OS cDNA library by PCR and ligated into the HindIII and BamHI sites of mEGFP-C1 construct (Addgene #54759) using ClonExpress II One Step Cloning kit (Vazyme, C112). Lentiviral plasmids (pLenti-TetOn-GFP-PARP10-T2A-Puro) were generated by cloning the human cDNA encoding PARP10 downstream of GFP in the construct digested with EcoRI and BamHI. The DNA fragments of human PARG103, ARH3, TARG1, MACROD2, DTX2, DTX3, DTX3L, and DTX4 were amplified from the cDNA library by PCR and ligated into the EcoRI and BamHI sites of the pLenti-CMV-mCherry construct. Mouse PARP10 was amplified from the cDNA library by PCR and ligated into the EcoRI and BamHI sites of pLenti-CMV-mEGFP construct. The DNA fragments of HA-Ub and Flag-Ub were synthesized by company (GeneUniveral) and cloned into the pcDNA3.1(+) (Invitrogen, V790-20) plasmid, digested with NheI and HindIII. Deletion mutants (ΔRRM2/3, ΔKH1, ΔKH2, ΔRRM1, ΔUIMs, ΔCD) and point mutations (G888W, H887E, H887A, 3RE, 3RA, 3RK, 3EA) of PARP10, PARG103 (PARG103-CD: E755A/E756A), ARH3 (ARH3-CD: D314E/T317A) DTX2 (DTX2 5 Mut: W24A, W114A, S568A, H582A, H594A) and mouse PARP10 (G837W, HRR>3E, ΔUIM) were created using Mut Express II Fast Mutagenesis kit (Vazyme, C214).

### CRISPR-Cas9 knockout and stable expression cell line

Guide RNAs targeting human PARP10 (5’-CAGGGTGGCCAGCAGTTCT-3’) was selected from Sanger Whole Genome knockout (KO) Arrayed CRISPR Libraries (Sigma Aldrich). U2OS cells were transiently transfected with U6-gRNA-RFP and pCMV-Cas9-BFP vectors together. The RFP/BFP-positive cells were sorted and single-cell cloning in a 96-well plate using a FACS cell sorter (Sony, MA900). The single-cell clones were confirmed for PARP10 deletion by western blotting using both commercial and homemade antibodies.

The selected gRNA^66^ (5’-CTGTGAGCTAGTTGATCTCG-3’) for MAVS was cloned into plasmid lentiCRISPR-V2 (Addgene #52961) digested by BsmBI. HEK29T cells were co-transfected with pVSV-G, psPAX2 and pLentiCRISPR-V2-MAVS-KO. The medium containing lentiviral particles was collected 48 hours post-transfection and filter-sterilized with a 0.45-μm filter. HEK293T cells were infected with lentiviral particles for 12 hours and subjected to puromycin (2 μg/mL) selection for a week. The KO pools were confirmed for MAVS deletion by western blotting.

To establish inducible TetOn-GFP-PARP10 WT/mutants cell lines, HEK293T cells were co-transfected with pVSV-G, psPAX2 and pLenti-TetOn plasmids (pLenti-TetOn-GFP-PARP10-WT/mutants-T2A-puro). To generate the lentivirus of reverse tetracycline-controlled transactivator (rtTA) for controlling the inducible expression, HEK293T cells were co-transfected pVSV-G, psPAX2 and pLenti-rtTA. The medium containing lentiviral particles was collected at 48 hours post-infection and filter-sterilized with a 0.45-μm filter. PARP10 KO or WT U2OS were infected with lentiviral particles of pLenti-TetOn and pLenti-rtTA for 12 hours. 2 μg/mL doxycycline was added for 12 hours to induce the expression of GFP-PARP10 WT/mutants. 2 μg/mL puromycin for 24 hours was used for the selection of GFP-positive cells.

### Immunofluorescence and confocal microscopy

Cells were seeded on 12-mm circular coverslips in 24-well culture plate for at least 12 hours. Cells were subjected to transfection and various treatments. Cells were fixed with 4% paraformaldehyde (Leagene, DF0135) for 10 min at room temperature, followed by permeabilization with 0.2% Triton X-100 (Sigma-Aldrich, T8787) in PBS for 10 min, and then blocked with 1% BSA (Genview, FA016-25G) in PBST (0.1% Tween-20 in PBS) for 30 min. The fixed cells were incubated with primary antibody in the blocking buffer overnight at 4 °C. After washing with PBST for three times, cells were incubated with Alexa Flour 488/555/647 conjugated secondary antibodies for 1 h at room temperature. After washing with PBST for three times, cells were mounted using ProLong™ Diamond Antifade Mountant with DAPI (Invitrogen, P36971). The cells were examined with Olympus FV3000 confocal laser scanning microscope equipped with a 60x oil objective.

### Threshold concentration analysis

TetOn-GFP-PARP10-WT/G888W/1-817(ΔCD) stable cells were seeded on 15 mm glass-bottom cell-culture dish. 2 μg/mL doxycycline (Solarbio, D8960) for 16 h and 0.02 μg/mL doxycycline for 12 h were used to induce the expression of TetOn-GFP-PARP10-WT and G888W/ΔCD, respectively. Under these conditions, nearly half of the cells showed condensates. Images of live cells were captured using a Nikon Spinning-Disk Confocal Microscope with a TIRF module, a live-cell imaging chamber, and a 60x oil-immersion objective. Raw files were analyzed using Imaris Workstation. A 2-μm diameter circular area was selected to calculate the mean intensity. The fluorescence intensity in the cytoplasm of cells that are dispersed or forming condensates is analyzed. The presence or absence of condensates was manually annotated. Cells with at least 3 foci are regarded as condensate-positive. The protein concentration was calibrated using GFP protein expressed i*n vitro* with a serial dilution of GFP concentration ranging from 0 to 16 μM in 50 mM Tris-HCl (pH 7.4), 150 mM NaCl buffer. The fluorescence intensity at the center of the GFP sample was measured and used to generate the standard curve. The cellular threshold concentration was calculated by converting the fluorescence intensity of the recombinant GFP protein concentration using the standard curve. At least 100 cells are measured and analyzed. Briefly, all the cells were binned based on concentration distribution using a square root number rule. The threshold concentration of certain proteins was defined as the average concentration of cells with >50% of cells forming condensates.

### Partition coefficient analysis

TetOn-GFP-PARP10-WT/G888W/ΔCD stable cells were grown on 15 mm glass-bottom cell-culture dish. The protein expression was induced with 2 μg/mL doxycycline for 48 h. Images of live cells were captured using a Nikon Spinning-Disk Confocal Microscope with a TIRF module, a live-cell imaging chamber, and a 60x oil-immersion objective. A 2-μm diameter circular area was selected to calculate the mean intensity. The ratio of fluorescence intensity between inside and outside the droplets was defined as the participation coefficient. The experiment was performed in triplicate, with each replicate having at least *n* = 50 cells.

### Fluorescence recovery after photobleaching (FRAP)

The fluorescence recovery after photobleaching (FRAP) experiment was performed using a Nikon Spinning-Disk Confocal Microscope with a TIRF module, a live-cell imaging chamber, and a 60x oil-immersion objective. TetOn-GFP-PARP10 WT/G888W/H887E cell lines were seeded on a 15-mm glass-bottom cell-culture dish. 2 μg/mL doxycycline for 48 h was used to induce the expression of GFP-PARP10 WT/G888W/H887E. The GFP-PARP10-WT cells were transiently transfected with mCherry-eAf1521 plasmids using PEI reagent. Time-lapse images were acquired using 20% intensity of 488/561 nm with 400-ms exposure. The condensates in a 2-μm diameter circular area were fully photobleached with 405 nm lasers at an intensity of 50% for 2 ms. The recovery process was recorded over a 60-second time course with 1-second interval after bleaching. The images were analyzed using Imaris Workstation. The mean fluorescence intensities from the area of background, photobleached, and non-photobleached cells were calculated. The background signals were subtracted from both photobleached and non-photobleached signals. The corrected fluorescence intensity over the time course was normalized by the pre-photobleached intensity and fitted to a curve using the LOWESS model in GraphPad Prism.

### Protein purification and antibody generation

Human recombinant PARP10 WT and mutants, including G888W, H887E, 3RE, and mouse PARP10 WT, were expressed in *E. coli* strain Rosetta (DE3) using the pET32a construct modified by adding a TEV protease recognition and cleavage site. Bacterial cells were grown to OD_600_ of 0.8 and induced with 1 mM IPTG (JSENB, JS0154) at 16 °C overnight. Bacteria were collected by centrifugation at 8,000 x *g* for 10 min and lysed by sonication in buffer containing 50 mM Tris-HCl (pH 7.4), 400 mM NaCl, 30 mM imidazole (Sigma-Aldrich, I2399-100G), 1xPMSF (Alpha Diagnostic, PMSF16-S-250), and 2 mM DTT (Sigma-Aldrich, 43815). After centrifugation at 18,000 x *g* for 20 min, the clear lysate supernatant was loaded onto a gravity Ni-NTA column (GenScript, L00666). After washing with lysis buffer, protein was eluted in 50 mM Tris-HCl (pH 7.4), 400 mM NaCl, 300 mM imidazole, 2 mM DTT. The protein was treated with TEV protease at 4 °C overnight. The cleaved protein was then purified using a Superdex 200 16/200 column (Cytiva, 28989335), equilibrated in SEC buffer containing 50 mM Tris-HCl (pH 7.4), 400 mM NaCl, and 2 mM DTT. Proteins were concentrated with Amicon Ultra centrifugal filters with a molecular weight cutoff (MWCO) of 10,000 (Millipore, UFC8010), snap-frozen in aliquots, and stored at −80 °C.

The purified human PARP10 WT and mouse PARP10 WT were used as the immunogen to immunize two New Zealand White rabbits via HUABIO (Hangzhou, China). The antisera of PARP10 were tested by western blot preliminarily, and the antibody of PARP10 was further subtracted from the antisera by the protein A/G purification.

Human PARP14 macrodomain-2-3-rFC, eAf1521-rFC, VSV-N-VHH1004 were synthesized by company (GeneUniveral) and cloned into pGEX-6P-1 vector. The sequence of VSV-N-VHH1004 single domain antibody was reported previously^67^. The constructs were expressed in *E. coli* strain Rosetta (DE3). Protein expression was induced with 1 mM IPTG at 16 °C overnight. Bacteria were collected by centrifugation at 8,000 x *g* for 10 min and lysed by sonication in the buffer containing 50 mM Tris-HCl (pH 7.4), 400 mM NaCl, 1xPMSF, and 2 mM DTT. After centrifugation at 18,000 x *g* for 20 min, the clear lysate supernatant was loaded onto a gravity column with glutathione resin. After washing with lysis buffer, the protein was eluted in 50 mM Tris-HCl (pH 7.4), 400 mM NaCl, 10 mM reduced glutathione, and 2 mM DTT. The eluted protein was then purified using a Superdex 200 16/200 column, equilibrated in SEC buffer containing 50 mM Tris-HCl (pH 7.4), 400 mM NaCl, and 2 mM DTT. Proteins were stored at 50% (v/v) glycerol supplemented with 0.02% sodium azide at −20 °C.

### Cryo-EM specimen preparation and data processing

0.45 mg/mL purified PARP10 was supplemented with 0.05% n-octyl-β-D-glucoside (Anatrace, O311) before being applied onto glow-discharged holey carbon grids (Quantifoil Au 300 mesh, R1.2/1.3). The grids were blotted for 3.5 s at 100% humidity using a Vitrobot Mark IV System (Thermo Fisher Scientific) before plunging into liquid ethane. The grids were screened on a Glacios Cryo-TEM (Thermo Fisher Scientific). A total of 12,772 movies were collected on Titan Krios G4 Cryo-TEM (Thermo Fisher Scientific) operating at 300 kV equipped with a Falcon 4i Direct Electron Detector and Selectris X Imaging Filter at a nominal magnification of 130,000× (corresponding to 0.92 Å/pixel) and an accumulated dose of 48 e^−^/Å^2^. Patch motion correction, patch CTF estimation, and the following processing steps were performed in CryoSPARC^68^. Micrographs with better than 10 Å resolution and relative ice thickness ranging from 1 to 1.2 were selected for particle picking. Particles were initially picked up with Topaz using a pretrained model^69^. Class averages selected from the second round of 2D classification were used as templates for training the Topaz model. The generated model was then used to select particles for the entire dataset again. Particles were sorted by 2D classification, and particles with clear structural features were chosen for generating initial maps. After several cycles of *ab initio* reconstruction and heterogeneous refinement, particles from the best class of heterogeneous refinement were applied to non-uniform refinement. The particles in the 3D class containing the highest copies of PARP10 were subjected to 3D variability analysis, and the generated particles were applied to non-uniform refinement.

### Structural analysis

AlphaFold3 was used to predict the structure of monomeric and oligomeric PARP10^70^. The predicted models were docked into the cryo-EM map of PARP10 in UCSF Chimera X^71,72^. To generate the model of PARP10 hexamer, six predicted models were docked into the cryo-EM map with the “Fit in Map” tool. Data collection and processing statistics are shown in Supplementary Table 1. Representative cryo-EM images, 2D class averages, and 3D maps are shown in Fig. 4b-e and Extended Data Fig. 4d,e. Structural analysis was performed, and structural images were generated in UCSF ChimeraX and PyMOL (Schrodinger).

### *In vitro* phase separation

The samples of *in vitro* phase separation were prepared by mixing the indicated concentration of recombinant protein with the crowding agent, including Ficoll-400 (Sigma-Aldrich, F2637) or PEG-8000 (Sigma-Aldrich, 89510), in a buffer containing 50 mM Tris-HCl (pH 7.4) and a determined concentration of NaCl. The samples were transferred to a sandwiched chamber generated by a cover glass and a glass slide with a double-sided spacer. Phase separation was observed using the DIC module of an Olympus wide-field microscope equipped with a 60x oil objective. All *in vitro* phase separation experiments were performed at room temperature, unless otherwise specified.

### *In vitro* ADP-ribosylation assay

The routine ADP-ribosylation assay was performed in a 50 μL reaction system containing 500 ng of enzyme, 1 mM NAD^+^ (MCE, HY-B0445), 50 mM Tris-HCl (pH 7.4), and 4 mM MgCl_2_. The reaction was conducted at 30 °C for 30 min and stopped by the addition of SDS sample buffer. For the experiment testing the effects of ADP-ribosylation on phase separation *in vitro*, 100 μM recombinant PARP10-WT or H887E protein was incubated with 12.5 mM NAD^+^ in reaction buffer at room temperature for the indicated times, followed by the addition of 10% Ficoll-400 to induce phase separation. For the experiment testing the roles of condensation on enzyme activity, 10 μM PARP10 WT protein was mixed with 10% (final concentration) PEG-8000 to generate condensates. 0.5 mM NAD^+^ was then added to initiate the reaction. The reactions were carried out at room temperature for the indicated times.

### Co-IP assays and TUBE pull down

HEK293T cells were transfected with indicated plasmid constructs. After 24 h of transfection, cells were collected and lysed on ice with lysis buffer (25 mM Tris-HCl, pH 7.4, 150 mM NaCl, 0.5% Triton X-100, 2 mM EDTA) supplemented with protease inhibitor cocktail (MCE, HY-K0010). 10 μM Olaparib was included in lysis buffer to prevent the artificial protein PARylation activated by DNA shearing *in vitro* during lysate preparation. Cell lysates were centrifuged at 4 °C for 10 min at 21,000 x *g*. The supernatants were incubated with agarose beads conjugated with GST-tagged GFP or mCherry Nanobody at 4 °C overnight. Beads were washed with lysis buffer four times and eluted with SDS-containing sample buffer. The resulting samples were analyzed by Western blotting. For the tandem ubiquitin-binding entities (TUBE) pull-down assay, we first created the GST-tagged 4xUBA construct by linking the human UBQLN1 UBA domain (536-589aa) with the GGGSGGG linker and purified the GST-fusion proteins. We followed the reported protocols for the pulldown^37,38^. In brief, cells were pre-treated with 10 μM proteasome inhibitor MG132 (MCE, HY-13259) for 4 h before sample collection. The lysates were prepared as described above. Cell supernatant was incubated with agarose beads conjugated with GST-4xUBA at 4 °C overnight. Beads were washed with lysis buffer four times and eluted with SDS-containing sample buffer. The resulting samples were analyzed by immunoblotting.

### Western blotting

Samples were boiled with 1x SDS sample buffer at 95 °C for 10 min, and then separated in 4%-20% Bis-Tris precast protein gels (ACE Biotechnology). Proteins were transferred to nitrocellulose membranes using the eBlot™ L1 Fast Wet Transfer System (GenScript). Membranes were blocked with 5% non-fat milk for 30 min at room temperature and then incubated with primary antibodies for 2 h at room temperature or at 4 °C overnight. After washing three times with TBST (0.1% Tween-20 in Tris-buffered saline), membranes were further incubated with IRDye goat-anti-rabbit IgG or goat-anti-mouse IgG secondary antibody for 1 h at room temperature. Membranes were washed three times with TBST, and then imaged with ChemiDoc MP Imaging System (Bio-Rad) or Odyssey CLx imager (LI-COR Biosciences). The quantification of immunoblots was performed by Image J (Open source Java program from NIH).

### CHX chase assay

TetOn-GFP-PARP10 WT stable cells were supplemented with 2 μg/ml doxycycline (Solarbio, D8960) for 24 h. 50 μg/ml CHX (MCE, HY-12320) was then added for the indicated time points, with or without treatment of 10 μM MG132 (MCE, HY-13259) or 0.5 μg/mL Bafilomycin A1 (MCE, HY-100558). Cells were lysed with 1x SDS sample buffer and subjected to western blotting.

### RT-qPCR

Total RNA was isolated with TRIzol^TM^ (Thermo Fisher Scientific, 15596018). RNA was reverse-transcribed into cDNA using the HiScript III 1st Strand cDNA Synthesis Kit (+gDNA wiper) (Vazyme, R312). qPCR was performed with technical duplicates using ChamQ Universal SYBR qPCR Master Mix (Vazyme, Q711-02) on CFX Connect Real-Time qPCR System (Bio-Rad). The quantification of target gene expression was normalized to the expression of housekeeping gene ACTB or GAPDH and was presented as fold changes relative to controls. Data are representative of at least three independent experiments.

### Virus binding, entry, and endocytosis assay

For virus binding assay, HEK293T cells transfected with the indicated constructs for 24 h were pre-cooled to 4 °C, followed by VSV infection (MOI=1.5) at 4 °C for 1 h. Cells were then washed with cool PBS for three times to remove the unbound VSV particles. For virus entry assay, cells were infected with VSV (MOI=1.5) at 4 °C for 1 h and transferred to 37 °C for 30 min to allow virus entry. Cells were then treated with 0.25% trypsin for 10 min at 4 °C to remove the VSV particles that had not entered, followed by three rinses with cold PBS. For virus endocytosis assay, cells were pre-treated with 10 μM Chloroquine 3 h before and during VSV infection (MOI=1.5, 6 hpi) at 37 °C. The VSV replication was quantified by RT-qPCR.

### Correlative light and electron microscopy (CLEM)

TetOn-GFP-PARP10 WT stable cells were seeded on gridded glass bottom dish (ibidi, 81166). To induce protein expression, 2 μg/mL doxycycline was added for 48 h. Cells were fixed with 4% paraformaldehyde for 20 min at room temperature. Fluorescence images were captured by Olympus FV3000 inverted confocal microscope. The cell morphology and the position were acquired and recorded under bright-field mode. Then, cells were fixed with 2.5% glutaraldehyde (Electron Microscopy Sciences, 16537-17) combined with 2% PFA in 0.1 M phosphate buffer PB (0.02 M NaH_2_PO_4_, 0.08 M Na_2_HPO_4_, PH 7.4) for 1h at room temperature, followed by three 15-min rinses with 0.1 M sodium cacodylate/PB. The post-fixation was performed with 1% osmium tetroxide (OsO4) (Ted Pella, 18466) in 0.1 M sodium cacodylate/PB (Aladdin, S464785) on the ice for 1 h, followed by three 10-min rinses with 0.1 M sodium cacodylate/PB and three washes with distilled water. Cells were then stained with 1% uranyl acetate (Electron Microscopy Sciences, 201119-31) in distilled water for 1 h at room temperature. After three washes with distilled water, cells were dehydrated in a cold-graded ethanol series (30%, 50%, 70%, 95%, and 100%, 10 min for each, 100% step repeated three times). Penetration was conducted with EPON 812 resin (Electron Microscopy Sciences, 14900) in 2:1 ethanol/resin for 30 min and 1:2 ethanol/resin for 30 min at room temperature, then in pure resin overnight, followed by three renewals with fresh resin for 3 h each. Samples were polymerized at 60 °C for 48 h, sliced into 80-nm-thick sections, and stained with uranyl acetate and lead citrate. Samples were examined under 120 kV Transmission Electron Microscope (Thermo Fisher Scientific, Talos L120C G2).

### MS-based metabolomics and metabolite assay

U2OS, BMDM, and HEK293T were infected with VSV (MOI=1.5) for indicated time periods. HEK293T transfected with GFP/GFP-PARP10-WT/GFP-PARP10-G888W were left untreated or infected with VSV (MOI=1.5) for 8 h. Each condition was performed in triplicate. Metabolites were extracted with 80% LC-MS grade pre-cold methanol (Thermo Fisher Scientific, A456-1) on dry ice. Cell pellets were separated by centrifugation at 16,000 x *g* for 15 min at 4 °C and supernatants were collected and dried by vacuum centrifugal concentrator (Martin Christ). The extracted metabolites were re-dissolved in 100 μL of 80% methanol for further analysis. The analysis was conducted using an LC-MS system comprising an Agilent 1290 Infinity II UHPLC system coupled with Agilent 6545 Q-TOF/MS. Chromatographic separation was performed on an ACQUITY UPLC BEH Amide column (100 mm×2.1 mm, 1.7 μm). The column was maintained at a flow rate of 0.3 mL/min and the injected sample volume was 5 μL. The mobile phase consisted of solvent A (15 mM ammonium acetate, 0.3% NH3·H20 in water) and solvent B (15 mM ammonium acetate, 0.3% NH3·H20 in 9:1 acetonitrile/water). The column was eluted with 95% solvent B for 1 min, followed by a linear gradient to 50% solvent B over 8 min, held at 50% for 3 min, a linear gradient to 95% solvent B over 0.5 min, and finally 1.5 min at 95% solvent B. The mass spectrometry data were obtained using electrospray ionization in both positive and negative ion mode over 50-1250 m/z. Raw data were processed using Profinder 10.0 (Agilent) for peak detection, alignment and integration. Throughout the analysis, QC RSD lower than 30% was selected. The metabolite levels were quantified based on the peak area and normalized to the protein contents determined by BCA assay. The enriched significant metabolites with *p* < 0.05 were used for metabolism pathway analysis with MetaboAnalyst 6.0 (https://dev.metaboanalyst.ca/ModuleView.xhtml). The top 10 enriched pathways and the included metabolites were chosen for plotting.

### NAD^+^ measurement

The NAD^+^ levels in cells were measured by ultra-performance liquid chromatography-mass spectroscopy as described in the metabolomics section or by bioluminescent assays using NAD/NADH-Glo™ kit (Promega, G9071) according to the manufacturer’s protocols. 50 μL of samples were mixed with 50 μL NAD/NADH-Glo™ Detection Reagent and incubated for 40 min in the 96-well plate. Then, the 96-well plates were read in the Varioskan LUX microplate reader (Thermo Fisher Scientific) with an integration time of 1 second per well. The *in vitro* reaction was carried out in a 20 μL system containing 10 μM recombinant PARP10 protein, with or without 10% PEG-8000, 200 μM NAD^+^, 150 mM NaCl, 50 mM Tris-HCl (pH 7.4) and 4 mM MgCl_2_. The reaction was conducted at room temperature for indicated times. The 20 μL reaction mixtures were incubated with 20 μL NAD/NADH-Glo™ Detection Reagent for the NAD^+^ measurement.

### RNA-sequencing and data analysis

BMDM were infected with VSV (MOI=1.5) for 12 h. HEK293T cells were transiently transfected with GFP vector or GFP-PARP10 WT/G888W/3RE for 24 h. Each condition was performed in triplicate. The total RNA from cells was extracted using TRIzol™ reagent (Thermo Fisher Scientific, 15596018), followed by the removal of genomic DNA with DNase I (NEB, M0303L). The RNA quality was verified with Nanodrop^TM^ OneC spectrophotometer (Thermo Fisher Scientific) for purity and LabChip GX Touch system (Revvity) for integrity. The total RNA was applied to library preparation using KCTM Digital mRNA Library Prep Kit (Seqhealth Tech. Co., Ltd., Wuhan, China) according to the manufacturer’s protocols. The kit utilized a unique molecular identifier (UMI) of 12 random bases for labeling cDNA molecules, which can eliminate the duplication bias and errors produced in the PCR and sequencing steps. The quantification and sequencing of enriched libraries with 200-500 bp fragments were performed by the DNBSEQ-T7 platform (MGI) with PE150 sequencing mode. RNA-sequencing reads were first filtered by fastp (v0.23.2) to remove low-quality reads and adaptor sequences. Clean reads were first clustered using UMI sequencing, followed by a new sub-clustering with > 95% sequence identity in the same cluster. For each sub-cluster, one consensus sequence was obtained by multiple sequence alignments. The processed sequences in BMDM and HEK293T were mapped to reference genomes GRCm38.102 and GRCh38.110, respectively, using STAR software (version 2.7.6a)^73^. The gene expression levels were calculated with FPKM (Fragments Per Kilobase of transcript per Million mapped reads) by normalizing gene counts from featureCounts (version 1.5.1). The averaged FPKM of three biological replicates were used for the comparison. Differentially expressed genes were identified using edgeR package (version 3.40.2). A *p*-value cutoff <0.05 and fold-change cutoff >2 was applied. Gene Set Enrichment Analysis (GSEA) was performed with GSEA software (version 4.4.0). The C2:curated gene sets from human MSigDB collections were chosen for analysis. The transcriptome data of whole blood samples from healthy and COVID-19 populations were obtained and analyzed in the COVID19db platform (http://www.biomedical-web.com/covid19db/home)^56^.

### Statistics

Statistical analysis was performed in GraphPad Prism. Data were presented as mean ± SD. The comparisons were performed by two-tailed unpaired Student’s *t*-test or one-way analysis of variance (ANOVA). A *p* value of less than 0.05 was considered significant. Statistical significance levels are denoted as follows: \**p* < 0.05, \*\**p* < 0.01, \*\*\**p* < 0.001, and \*\*\*\**p* < 0.0001.

## Data availability

The RNA sequencing data are available in Genome Sequence Archive (https://ngdc.cncb.ac.cn/gsa/) of China National Center for Bioinformation with identifier CRA027841 (Bulk RNA-seq data in BMDM), and in Genome Sequence Archive (https://ngdc.cncb.ac.cn/gsa-human/) with identifier HRA012338 (Bulk RNA-seq data in HEK293T). The metabolomic data are available in OMIX (https://ngdc.cncb.ac.cn/omix/) of China National Center for Bioinformation with identifier OMIX012645. The cryo-EM maps were deposited in the Electron Microscopy Data Bank with accession ID: EMD-65291.

## Supplementary information

**Supplementary Table 1**

Cryo-EM data collection and processing

**Supplementary Table 2**

Primers for RT-qPCR

**Supplementary Table 3**

Bulk RNA-seq results of VSV-infected BMDM

**Supplementary Table 4**

Bulk RNA-seq results of PARP10-expressing HEK293T

## Notes

### Competing Interest Statement

The authors have declared no competing interest.

